# Empirical estimates of the mutation rate for an alphabaculovirus

**DOI:** 10.1101/2021.09.07.459225

**Authors:** Dieke Boezen, Ghulam Ali, Manli Wang, Xi Wang, Wopke van der Werf, Just M. Vlak, Mark P. Zwart

**Affiliations:** Department of Microbial Ecology, The Netherlands Institute of Ecology (NIOO-KNAW), Wageningen, The Netherlands; Laboratory of Virology, Wageningen University and Research, Wageningen, The Netherlands; Wuhan Institute of Virology, Chinese Academy of Sciences, Wuhan, PR China; Centre for Crop Systems Analysis, Wageningen University and Research, Wageningen, The Netherlands

## Abstract

Mutation rates are of key importance for understanding evolutionary processes and predicting their outcomes. Empirical estimates of mutation rate are available for a number of RNA viruses, but few are available for DNA viruses, which tend to have larger genomes. Whilst some viruses have very high mutation rates, lower mutation rates are expected for viruses with large genomes to ensure genome integrity. Alphabaculoviruses are insect viruses with large genomes and often have high levels of polymorphism, suggesting high mutation rates despite evidence of proofreading activity by the replication machinery. Here, we report an empirical estimate of the mutation rate per base per strand copying (s/n/r) of Autographa californica multiple nucleopolyhedrovirus (AcMNPV). To avoid biases due to selection, we analyzed mutations that occurred in a stable, non-functional genomic insert after five serial passages in *Spodoptera exigua* larvae. Population bottlenecks, viral mode of replication and thresholds for mutation detection likely affect mutation rate estimates, and we therefore used population genetic models that account for these processes to infer the mutation rate. We estimated a mutation rate of 1×10^−7^ s/n/r. This estimate was not sensitive to different model assumptions or including whole genome data. The rates at which different classes of mutations accumulate provide good evidence for neutrality of mutations occurring within the inserted region. We therefore present a robust approach for mutation rate estimation for viruses with stable genomes, and strong evidence of a much lower alphabaculovirus mutation rate than supposed based on the high levels of polymorphism observed.

**Author Summary:** Virus populations can evolve rapidly, driven by the large number of mutations that occur during virus replication. It is challenging to measure mutation rates because selection will affect which mutations are observed: beneficial mutations are overrepresented in virus populations, while deleterious mutations are selected against and therefore underrepresented. Few mutation rates have been estimated for viruses with large DNA genomes, and there are no estimates for any insect virus. Here, we estimate the mutation rate for an alphabaculovirus, a virus that infects caterpillars and has a large, 134 kilobase pair DNA genome. To ensure that selection did not bias our estimate of mutation rate, we studied which mutations occurred in a large artificial region inserted into the virus genome, where mutations did not affect viral fitness. We deep sequenced evolved virus populations, and compared the distribution of observed mutants to predictions from a simulation model to estimate mutation rate. We found evidence for a relatively low mutation rate, of one mutation in every 10 million bases replicated. This estimate is in line with expectations for a virus with self-correcting replication machinery and a large genome.

## Introduction

Mutation rates are of key importance for understanding and predicting evolutionary patterns, as the mutation rate modulates the mutation supply of a population [1]. Large mutation supplies can fuel rapid and repeatable adaptation [2,3], but also increase the mutational load on a population [4]. By contrast, low mutation supplies can limit the rate of adaptation [5], but also result in a lower mutational load [4]. The impact of mutational supply also depends on the topography of the fitness landscape. Small mutational supplies can have advantages for evolution on rugged fitness landscapes: although adaptation will be slower and in most cases less fit genotypes will be selected, some populations can avoid becoming trapped on local fitness peaks [6]. Mutation rates are not only relevant to understanding basic evolutionary processes, but they also impinge on real world outcomes, such as the efficacy of prophylactic or therapeutic interventions to infectious diseases [7,8].

Viruses have high mutation rates [7,9,10], with estimates of mutations per site per strand copying (s/n/r) ranging from 2×10^−8^ for Enterobacteria phage T2 [11] to 2×10^−4^ for Influenza A virus [12]. Whilst these high mutation rates are thought to contribute to the rapid adaptation of viruses, the majority of mutations are typically deleterious [13,14]. Many viruses with large genomes belong to Group I of the Baltimore classification (dsDNA viruses) [15], and typically have polymerases with proofreading activity, which should enhance the fidelity of replication [16]. A general expectation is therefore that viruses with relatively large genomes have lower mutation rates [9]. An inverse relationship between genome size and mutation rate indeed has been found. Small genomes can tolerate higher mutation rates as a larger proportion of mutation-free genomes are generated in each round of replication, due to their small size [9,17].

The alphabaculoviruses are a large group of insect baculoviruses that have been studied because of their biocontrol and biotechnological potential [18,19]. Alphabaculoviruses have relatively large dsDNA genomes compared to other viruses [20], and high levels of within-host genetic diversity have been documented from wild [21–23] and captive [24] insect populations. It has been suggested that baculoviruses might therefore have high mutation rates despite their large genome sizes [24]. To our knowledge, no empirical estimates of the mutation rate have been reported for any baculovirus or insect virus to date, and there are only a few estimates for other large dsDNA viruses [9].

A major challenge for making empirical estimates of mutation rates is the need to account for biases due to selection [9]. Studies on the distribution of mutational fitness effects (DMFE) have shown that most mutations in viral genomes are deleterious [13,14], and selection will remove these mutations from populations. By contrast, beneficial mutations will increase in frequency due to selection. These opposing effects of purifying and directional selection make it problematic to derive information on mutation rates directly from mutation accumulation in a viral genome. Many different approaches have been developed to remove the bias introduced by selection [7,9]. For example, some studies considered the frequency of lethal mutations in a population, since these variants cannot replicate autonomously and therefore represent a snapshot of genetic variation [25]. Others have evolved viruses in hosts expressing a viral gene, and then restricting their analysis to the sequence of this redundant viral gene [26]. Another strategy reported recently has been to incorporate fluorescent markers with inactivating mutations into a viral genome, and then performing fluctuation tests based on recovering fluorescence [12].

In the current study, we report the first empirical estimate of mutation rate for a large dsDNA virus, the alphabaculovirus Autographa californica multiple nucleopolyhedrovirus (AcMNPV). To ensure selection did not bias our estimates, we analyzed virus populations carrying a large, nonfunctional genomic insert that was stably maintained [27], exploiting the genomic stability of the Group I viruses [28]. For our analysis, we assumed that mutations in this region are neutral due to the absence of known viral genes and regulatory sequences, and verified this assumption. We also developed a population genetics model to estimate mutation rates from empirical data, that incorporates the effects of population bottlenecks and different modes of virus replication on the occurrence and maintenance of mutations. Using this approach, we made robust estimates of the mutation rate for a baculovirus for the first time.

## Results and Discussion

### Serial passage and detection of mutations

To estimate the mutation rate of a large dsDNA virus, we experimentally evolved a variant of alphabaculovirus AcMNPV containing a stable, non-functional genomic region. The AcMNPV variant used was a so-called bacmid: an infectious clone that also contains the AcMNPV genome (~134 kb). It also contains bacterial sequences that enable propagation as a low copy number plasmid in *Escherichia coli* (~12.5 kb) and the acceptance of expression cassettes by transposition [27]. The specific variant used here contains an expression cassette from the pFastBac-Dual vector to restore expression of complete and functional polyhedrin (Figure 1). We consider the inserted bacterial sequences to be non-functional and therefore neutral in insects, except for the polyhedrin promoter and open reading frame (ORF) sequences derived from the pFastBac Dual vector. This renders an 11,646 bp neutral region for a mutation accumulation experiment (Figure 1). By contrast, the remainder of the AcMNPV genome is intact and unaltered, making this system representative for this virus. We will hereafter refer to this bacmid-derived AcMNPV variant with restored polyhedrin expression as “BAC”. The virus could be reconstituted from the infectious clone by transfection of the BAC genome into fourth instar (L4) *Spodoptera exigua* (Hübner) (see Materials and Methods). Mutations were allowed to accumulate across the BAC genome by experimentally evolving five replicate BAC lineages (referred to as lineages A, B, C, D and E) for five passages in *S. exigua* L2. For each replicate lineage, passaging was performed in five larvae exposed to a high viral dose of occlusion bodies (OBs) sufficient to kill all larvae. Upon death larval cadavers were collected and pooled prior to the isolation of OBs, which were used to inoculate larvae orally for the next passage. Further details on the experimental evolution experimental setup can be found in the Materials and Methods.

**Figure 1:**
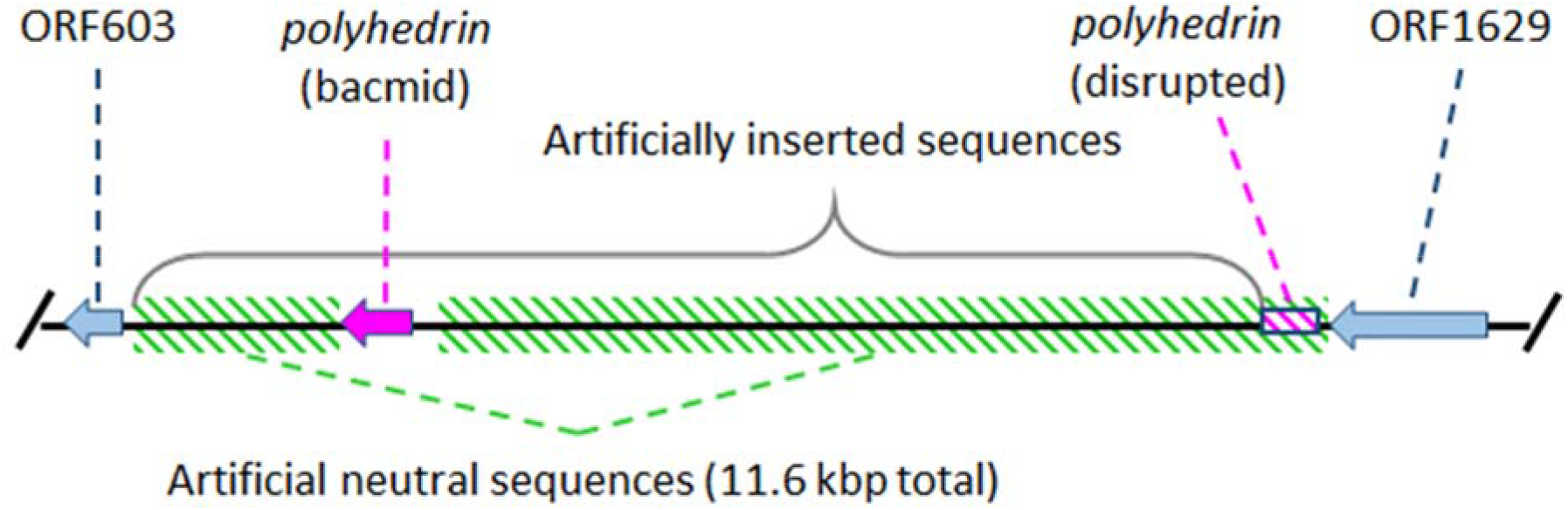
Illustration of the neutral region (green diagonal bars) used for estinating mutation rate. ORF603 and ORF1629 (light blue) are native AcMNPV genes, between which lies the *polyhedrin* gene which was disrupted (box with magenta bars) by the insertion of the bacmid sequences in its 5’ end. To restore *polyhedrin* expression the gene has been reinserted under control of its native promotor within the bacmid insert. We consider the sequences of bacterial origin in the insert and the remnants of the pseudogenized *polyhedrin* gene copy as neutral sequences.

Evolved lineages A - E, as well as the ancestral BAC were sequenced using Illumina HiSeq to detect mutations in both the non-functional genomic insert, as well as across the whole baculovirus genome. When mapping reads to the BAC reference genome we noticed a correlation between low sequencing coverage and the calling of mutations, and we therefore removed regions with relatively low coverage (Figures S1–S4, see Materials and Methods). We show that mutations indeed accumulated across lineages and passages (Table 1). When analyzing the sequence data, the number of mutations detected is dependent on the minimum threshold value (*τ*) for mutation frequency (e.g. *τ* values of 0.5, 1, 2% were used). Regardless of the chosen mutation frequency threshold, the number of mutations detected is low for both the bacmid insert and genome after five passages in *S. exigua* L2.

**Table 1:** Mutations called per lineage, where *τ* is the threshold frequency for detecting mutations.

| Lineage | Neutral region only |  |  | Whole genome |  |  |
| --- | --- | --- | --- | --- | --- | --- |
| | $\tau = 0.5\%$ | $\tau = 1.0\%$ | $\tau = 2.0\%$ | $\tau = 0.5\%$ | $\tau = 1.0\%$ | $\tau = 2.0\%$ |
| BAC | 1 | - | - | 1 | - | - |
| A | 2 | - | - | 20 | 7 | 1 |
| B | 2 | - | - | 14 | 3 | 2 |
| C | 1 | 1 | - | 9 | 2 | 1 |
| D | 1 | 1 | - | 7 | 6 | 2 |
| E | 2 | 1 | 1 | 18 | 2 | 2 |

### Low mutation rate for AcMNPV with bacmid and whole-genome mutation data

To make robust inferences on the mutation rate (*μ)* from these experimental data, we developed a population genetic model. Briefly, we generated a stochastic model simulating neutral evolution in a virus genome, modelled as mutation frequencies per base per viral generation. We fitted this model to the experimental data by considering the number of bases with a frequency of mutations above the threshold *τ*, using a maximum likelihood approach. We estimated the viral mutation rate to be ~1 × 10^−7^ s/n/r, with all estimates for the neutral bacmid region and the whole genome data being similar (Figure 2). Below we describe these results in more detail.

**Figure 2:**
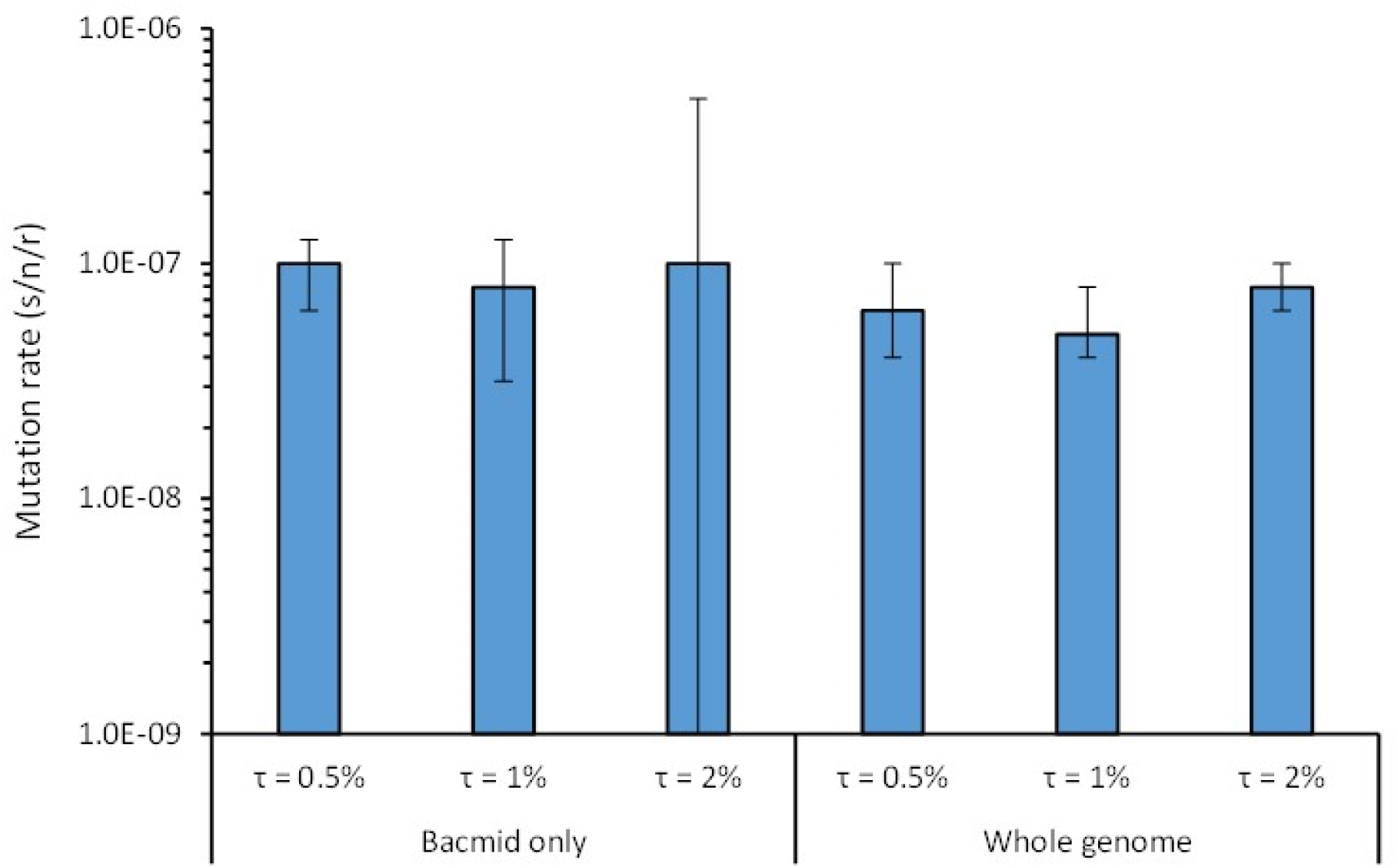
Mutation rate estimates (s/n/r, mutations per site per strand copying) derived with the model are given based on the neutral bacmid region or the whole genome. Different values for the threshold frequency for detecting mutations (*τ*) were used, noted as percentages here. Error bars represent the 95% fiducial limits, as determined by bootstrapping. Note that for the neutral region and *τ* = 2.0%, the lower fiducial limit extends to zero due to the low number of mutations detected.

The simulation model was run for a range of model parameter values for mutation rate, viral replication modes (*ρ* values of 1, 3 and 10; see also Materials and Methods section), threshold values for mutation detection (*τ* values of 0.5%, 1% and 2%), and lengths corresponding to the neutral bacmid region only and the whole virus genome (see Materials and Methods section for details). Strikingly, our mutation rate estimates were robust relative to the replication mode *ρ* value and to the choice of experimental data used (neutral region or whole genome), with estimates of approximately 1×10^−7^ s/n/r in all scenarios (Figure 2, Figure S5–S6). We also estimated the mutation rate with established models for sequencing data from clones [9], adapting these methods to use the frequency of mutations determined by high-throughput sequencing data instead of Sanger sequences from clones (see Materials and Methods). These estimates were similar to those obtained with the first approach, although they tended to be about a factor 2.5 smaller (*μ* ≤ 4 ×10^−8^ s/n/r, Figure S7). This difference was expected, as this alternative approach does not take into consideration the effects of the mutation detection threshold *τ*, but rather assumes all mutations will be detected irrespective of their frequency. As the sensitivity limits of the deep-sequencing data is not taken into account, this approach will underestimate the mutation rate.

One of the purported strengths of the first approach is that we have a large region (11.6 kb) in which point mutations will not affect fitness, given there are no known viral genes or regulatory sequences (Figure 1). Although large genomic deletions in this region would presumably be beneficial as they could speed up viral replication [28], none were detected: Sequence coverage was similar to the rest of the genome for the evolved populations (Figures S1–S4), ruling out the occurrence of large deletions at high frequencies. To test if point mutations in this region were indeed neutral, we considered the rate at which mutations occurred in the neutral intergenic region (dI) normalized by the rate of synonymous substitutions in viral genes (dS) (see Materials and Methods). If mutations in the inserted region are neutral, we expect dI/dS ~ 1. For the different values of *τ* used, dI/dS estimates were indeed approximately 1 (Figure 3), ranging from 0.72 to 1.96. For the condition under which we could perform a formal test (*τ* > 0.5%), dI/dS was not significantly different from 1 (One-sample *t-*test: *t*_4_ = 1.685, *p* = 0.167), providing support to the idea that mutations in the bacmid region are neutral. By contrast, when the same analysis was performed for the rate of nonsynonymous mutations (dN) in viral genes, all dN/dS values were ≤ 0.34 (Figure 3). For the condition under which we could perform a formal test (*τ* > 0.5%), dN/dS was significantly lower than 1 (*t*_4_ = −5.717, *p* = 0.005). This underrepresentation of nonsynonymous mutations in the viral genome presumably occurs because most nonsynonymous mutations will be deleterious [13,14], and therefore are removed by purifying selection. Despite evidence for purifying selection acting on viral genes, mutation rate estimates were similar for the neutral region and the whole genome (Figure 2).

**Figure 3:**
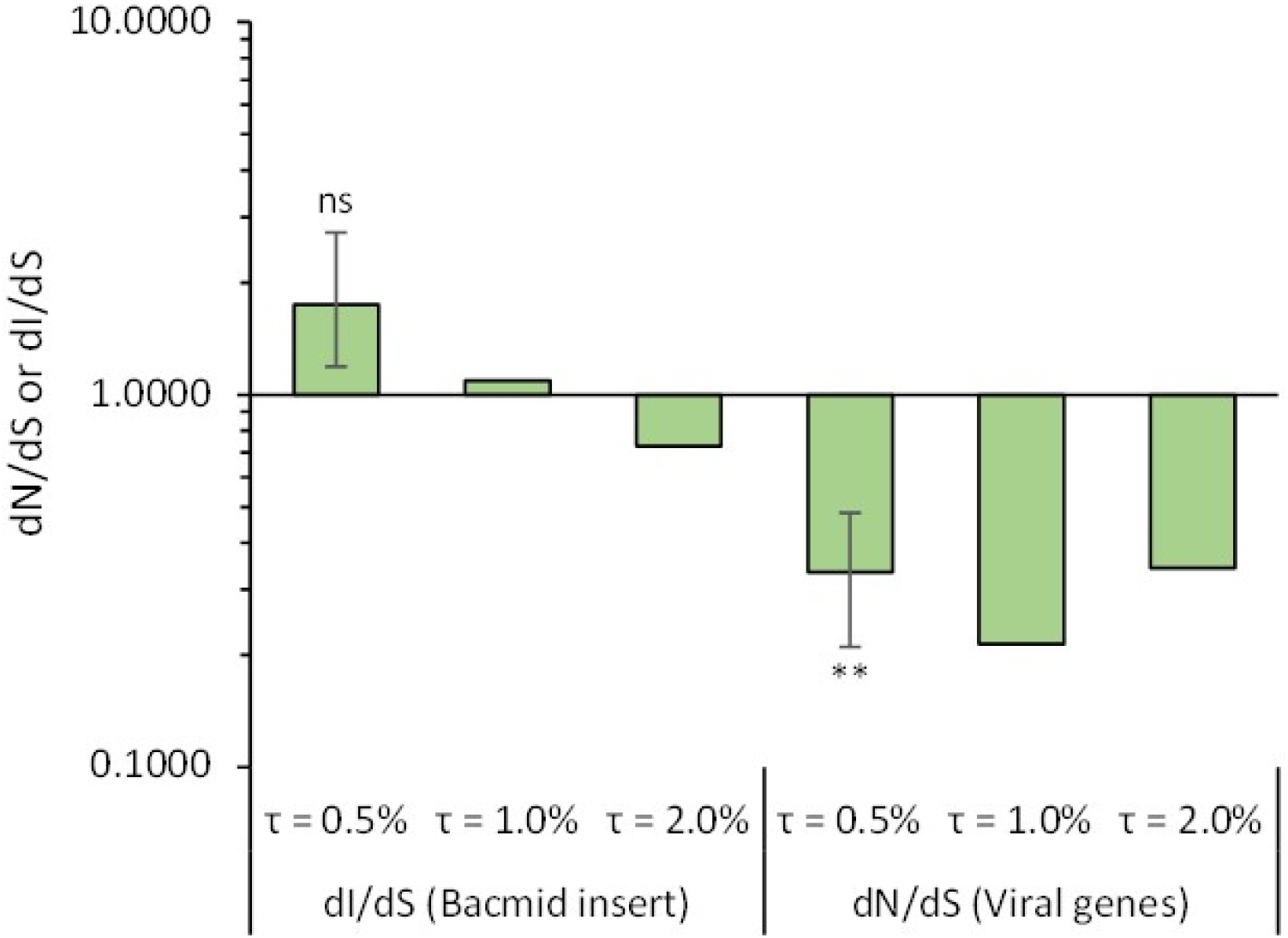
The rate of mutations in the artificial neutral regions (dI), normalized by the rate of synonymous mutations in native viral genes (dS), is shown by the three bars on the left. The values are close to 1, indicating neutral evolution. The three bars on the right show the rate of non-synonymous mutations in viral genes (dN), normalized by dS. These values are less than 1, indicating purifying selection. dI/dS and dN/dS values were determined for different values for the threshold frequency for detecting mutations (*τ*), noted as a percentage in the figure. Only for the lowest threshold value for mutation detection (*τ* = 0.5%) were there synonymous mutations detected in each sample, and therefore we could perform statistical analysis only for this condition. Error bars indicate the 95% confidence interval of the estimate as determined by bootstrapping, and results of a one-sample *t*-test are indicated by ns (non-significant, *p* > 0.05) and ** (*P* < 0.01).

The accumulation of mutations in a virus population is affected by the viral replication mode (*ρ*) [9,10,29–31], and the distribution of mutation frequencies is linked to the mode of replication [31]. Here we considered this effect by fitting the genome evolution model with values *ρ* = 1 (the “geometric growth” scenario), *ρ* = 3 (mixed replication scenario) and *ρ* = 10 (“stamping machine” scenario). We found that viral mode of replication had a consistent but small effect on the estimated mutation rate (Figure S6), with higher values of *ρ* corresponding to higher mutation rate estimates, as expected. The effect on model fit was minimal (Figure S5), and we therefore cannot make inferences on the mode of replication from these data. This result is not surprising, given that the number of mutations we detected was small and that the model fitting only considers the number of sites with a mutation frequency above *τ*, and not the frequency of individual mutations. For baculoviruses, the mode of replication has not been described formally, to the best of or our knowledge. However, as these viruses probably employ rolling circle amplification [32], replication is likely to follow the “stamping machine” scenario and to be described best by high values of *ρ*.

### AcMNPV mutation rate estimate is congruent with estimates for other viruses

Mutation rates (s/n/r) have been estimated for a number of viruses, allowing for a comparison with our baculovirus estimate. For this comparison, we included a collection of mutation rate data [9,33], with updated mutation rates for the RNA viruses influenza A virus [12] and poliovirus [31] due to the availability of better estimates. When multiple mutation rates were available for one virus, we used only the most recent estimate because methodological advances make estimates that are more recent more reliable. Our estimate clearly is congruent with mutation rate estimates for other dsDNA viruses, as it is close to the predicted relationship between genome size and mutation rate (Figure 4). As AcMNPV has a relatively large genome, it is also one of the lowest estimates of mutation rate in DNA viruses reported in the literature, similar to that of Escherichia virus *λ*, and with only Enterobacteria phage T2 (170 kbp) being lower.

**Figure 4:**
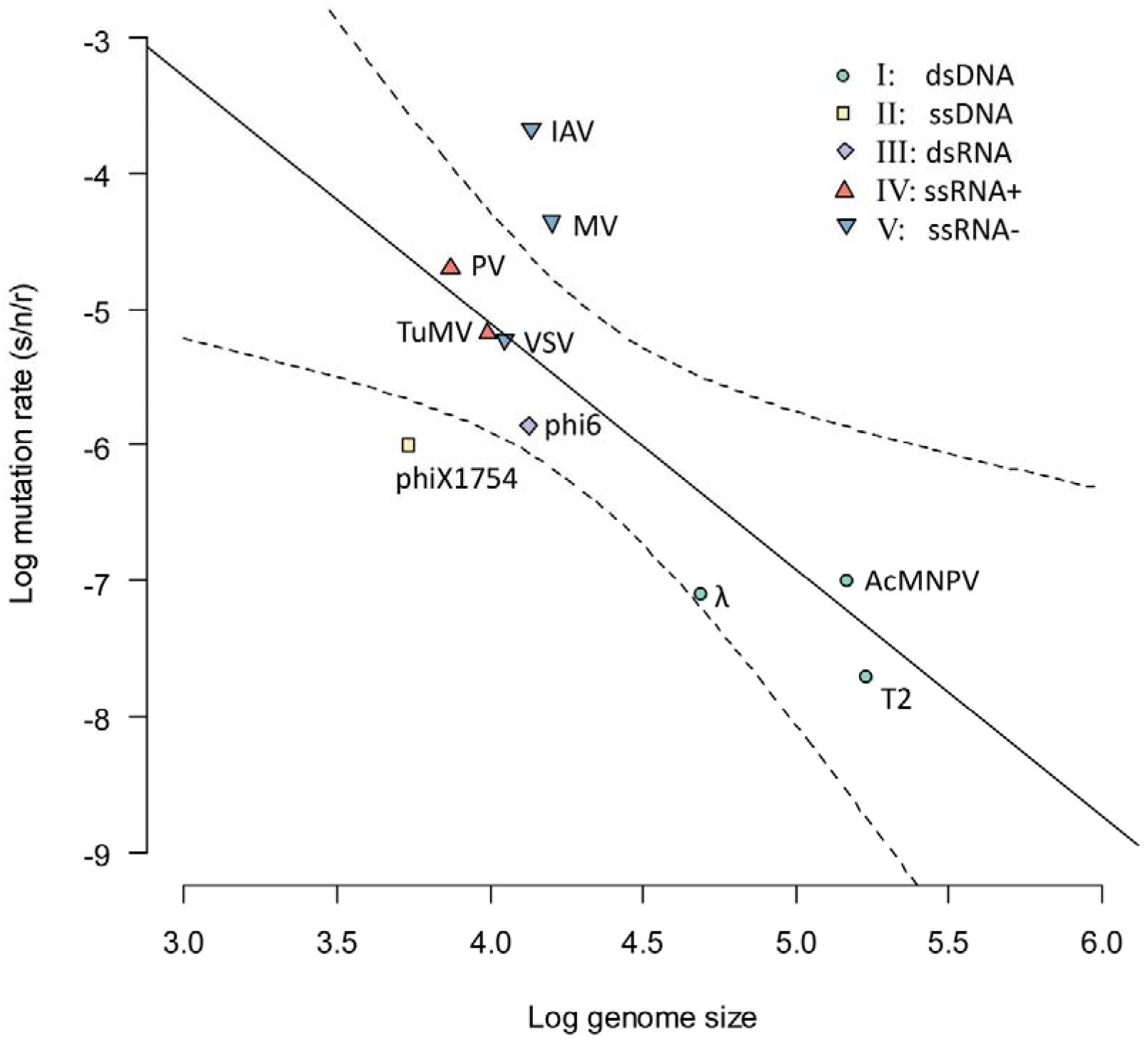
An overview of known mutation rate estimates (s/n/r, ordinate) for viruses with different genome sizes (abscissa) is given. The color and shapes of symbols indicate the Baltimore classification group to which each virus belongs, as indicated by the legend in the top right of the figure. The solid black line marks the regression line, with dotted lines marking the 95% confidence interval. The slope of the fitted relationship is significantly less than zero (*t*_8_ = −3.243, *p* = 0.012), and the coefficient of determination (*r*^2^) is 0.568. Our mutation rate estimate for AcMNPV is in good agreement with the fitted relationship between genome size and mutation rate. Other virus names in the figure are Enterobacteria phage T2 (T2), Escherichia virus λ (λ), Escherichia virus ΦX174 (phiX174), Influenza A virus (IAH), Measles virus (MV), Poliovirus (PV), Pseudomonas virus Φ6 (Phi6), Turnip mosaic virus (TuMV), and Vesicular stomatitis virus (VSV).

Besides being consistent with trends in other viruses, our estimate of mutation rate for AcMNPV is congruent with what is known about its polymerase. AcMNPV codes for a DNA-dependent DNA polymerase (DNApol) that belongs to the family B DNA polymerases and contains the exonuclease domain, thought to be responsible for proofreading and editing of mismatches [16,34,35]. The 3’ to 5’ exonuclease activity of this domain - essential for repairing errors - has been confirmed [36], and hence a low mutation rate is expected for AcMNPV. The other viruses with low mutation rates are both dsDNA bacteriophages with relatively large genomes (Figure 4). Proofreading activity also has been demonstrated in the case of T2 [37].

### Analysis of the mutational spectrum: a low transition to transversion ratio?

We considered whether our data shed light on AcMNPV’s mutation spectrum, the occurrence of the different kinds of single-nucleotide mutations, focusing on the transition to transversion ratio (Table 2; see also Table S1). For the neutral region, the number of observed mutations is too small to be informative, even for the lowest mutation detection threshold of *τ* = 0.5%. For the whole genome and for the lowest mutation detection threshold, transitions are close to the expected ratio in the absence of mutation bias (0.5), which may indicate a low transition to transversion ratio. However, this trend does not hold for higher values of *τ*, and selection may bias the mutations detected outside of the neutral region. Due to the small number of mutations detected in the neutral bacmid region and the possible effects of selection on the whole-genome data, we therefore cannot draw any firm conclusions on AcMNPV’s mutational spectrum. Analysis of a larger number of neutral mutational events will be necessary to draw conclusions on mutation spectrum. These larger numbers could be achieved by analyzing a larger number of replicate populations or populations evolved over a longer period of time, although improved sequencing methods with much lower error rates [38] would probably suffice to analyze mutation bias for the evolved populations described here.

**Table 2:** Overview mutation spectrum

| Observation or index | Relative frequency of mutation |  |  |  |
| --- | --- | --- | --- | --- |
|  | Whole genome |  |  | Bacmid only |
| | $\tau = 0.5$ | $\tau = 1.0$ | $\tau = 2.0$ | $\tau = 0.5$ |
| Transitions / Transversions | 28/50 | 9/11 | 6/2 | 4/5 |
| Ratio<sup>a</sup> [95% confidence interval] | 0.56 [0.34-0.91] | 0.82 [0.30-2.17] | 3.00 [0.54-30.3] | 0.80 [0.15-3.72] |

<sup>a</sup> The ratio of transitions to transversions

### Concluding remarks

Our low mutation rate estimate is congruent with the large genome size of AcMNPV and its known proofreading activity [36]. By contrast, the high genetic variation often observed within alphabaculovirus populations [21–24] remains a conundrum. In one case frequency-dependent selection has been observed and may account for stable polymorphisms [39,40]. Whether intricate relationships between genetic variants that complement each other are common or evolutionarily stable remains to be seen, but such relationships would help to explain the genetic diversity often seen within alphabaculovirus populations.

We estimated the mutation rate for the alphabaculovirus AcMNPV, using a novel approach that depends on the insertion of an artificial neutral region in the viral genome, and which starts with a single genotype and exploits the genome stability of group I dsDNA viruses. Such an approach requires mutations in this ‘artificial’ region to be neutral, and dI/dS results suggest this assumption is met. Whereas others have employed rolling circle amplification of sequencing templates to reduce sequencing error [38], here we developed an approach to analyzing regular Illumina sequencing data that were not gathered specifically with the intent of determining mutation rates. Our approach relies on a detailed analysis of the sequencing data to eliminate obvious sequencing biases and a comparison of sequencing data to simulation-model predictions. The use of models allows us to take into consideration the effects of viral demography on mutation rate estimates, and also limit the impact of choosing thresholds for mutation detection, as the same threshold is applied in the model.

## Materials and Methods

### Experimental evolution of AcMNPV

pBac-E2 (BAC) was evolved experimentally in *S. exigua* larvae in five replicate lineages (A, B, C, D and E). To reconstitute the virus from the bacmid, the haemocoel of *S. exigua* L4 was injected with a total volume of 20 μl (i.e., 2 × 10 μl), containing a 4:1:1 mixture of Lipofectin transfection reagent (ThermoFisher Scientific), water and BAC DNA (~ 15 μg DNA per larva). Upon larval death, OBs were harvested from cadavers. From a single infected larva, OBs were isolated and diluted to 2×10^7^ OBs/ml. Serial passage of BAC was performed five times for each replicate lineage, with five larvae used for each replicate lineage. For each passage, newly molted L2 were starved for 12 h and then inoculated by droplet feeding with an OB suspension exceeding 10 x LC_99_ (≥ 2×10^7^ OBs/ml), to avoid narrow transmission bottlenecks. Per replicate lineage, five inoculated larvae were transferred to 6-well tissue culture plates with artificial diet plugs. Larvae were incubated at 26°C and with a 14 h:10 h day-night photoperiod. Upon death, larval cadavers were collected, pooled and used to inoculate the next passage. After five passages, lineages A - E were amplified using 100 *S. exigua* L3, and 1.5 × 10^9^ OBs were used to extract viral genomic DNA. Briefly OBs were dissolved with DAS buffer (0.1 M Na_2_CO_3_, 0.15 M NaCl, 10 mM EDTA, pH 11), and DNA was then extracted from the liberated occlusion-derived virus particles using a DNA isolation kit (Omega Bio-tek) following the manufacturer’s instructions. For the BAC, DNA was extracted from 50 ml LB from an overnight culture using a plasmid midi kit (Qiagen). Successful genomic DNA extraction was confirmed by PCR with primers gp41 inner F (5’-CAAGAGCAAAGAACCGACG-3’) and inner R (5’-TTATGCAGTGCGCCCTTTCGT-3’), and contamination of SeMNPV was ruled out by PCR with primers Se F (5’-GACGACGAATTATGTTGTGACCGAC-3’) and R (5’-AGATGGATGGAAAGGCAACGCT-3’). Purified DNA (~ 1.5 ug) from the evolved AcMNPV lineages A, B, C, D and E, as well as the ancestral BAC, was sequenced by Illumina HiSeq PE150 (Beijing Novogene Bioinformatics Technology Co., Ltd).

### Mutation calling and filtering

Because the number of viral reads was not equal across samples, fastq files were subsampled to ensure an approximately equal mean coverage across the reference genome for each isolate using seqtk sample [41]. NGS data was analyzed using CLC Genomics Workbench 20.0 [42]. Reads were trimmed (quality limit = 0.05) and mapped to the reference genome (see Online Materials for sequence and annotation files). Mutations were called using the “low frequency variant detection tool” (minimum frequency = 0.5%). An overview of parameter settings can be found in the Supplementary Online Materials (Appendix 1).

Due to the small likelihood of the same mutation occurring independently in two or more populations, we assume mutations found in two or more evolved populations were present in the ancestral population (having been present at low frequency in the BAC DNA or occurring *de novo* after reconstitution of the virus. Therefore, mutations called in multiple lineages and/or in the ancestral BAC isolate were filtered out. Additional filtering criteria were: forward-reverse balance > 0.05, read count > 10, and the type of mutation is “SNV” (single nucleotide variant). Additionally, positions with extreme coverage values were excluded. To this end, we ranked the coverage value per position for each lineage and excluded the upper and lower 1%. Analyses were done with three thresholds (*τ*) for mutation frequency: 0.5%, 1% and 2% (see Supplementary Online materials, Appendices 2-4). Finally, mutations were tallied per isolate, both across the whole genome and for the neutral bacmid insert.

### Mutation model and mutation rate estimation

We generated a stochastic model that predicts the distribution of mutation frequencies per base in an evolving virus genome, and then fitted this model to our empirical data with a maximum likelihood approach to obtain mutation rate estimates. The model was implemented in R 4.0.3 [43] and all code is available (see Supplementary Online Material, Appendices 5-6). We model the genome region under consideration as a vector with *g* elements for each nucleotide position, with each element representing the total frequency *f* of mutated bases at position *i*. For simplicity, we do not consider the identity of the mutated bases and we do not allow for reversions, as we are considering scenarios in which mutations are rare and the probability of a reversion occurring and reaching high frequency is very low. We assume that all mutations are strictly neutral, and that all changes in mutation frequency result from the occurrence of *de novo* mutations or neutral processes like stochastic changes in allele frequencies due to population bottlenecks (i.e. genetic drift). All model parameter values are given in Table 3, together with additional explanation and justification.

**Table 3:**
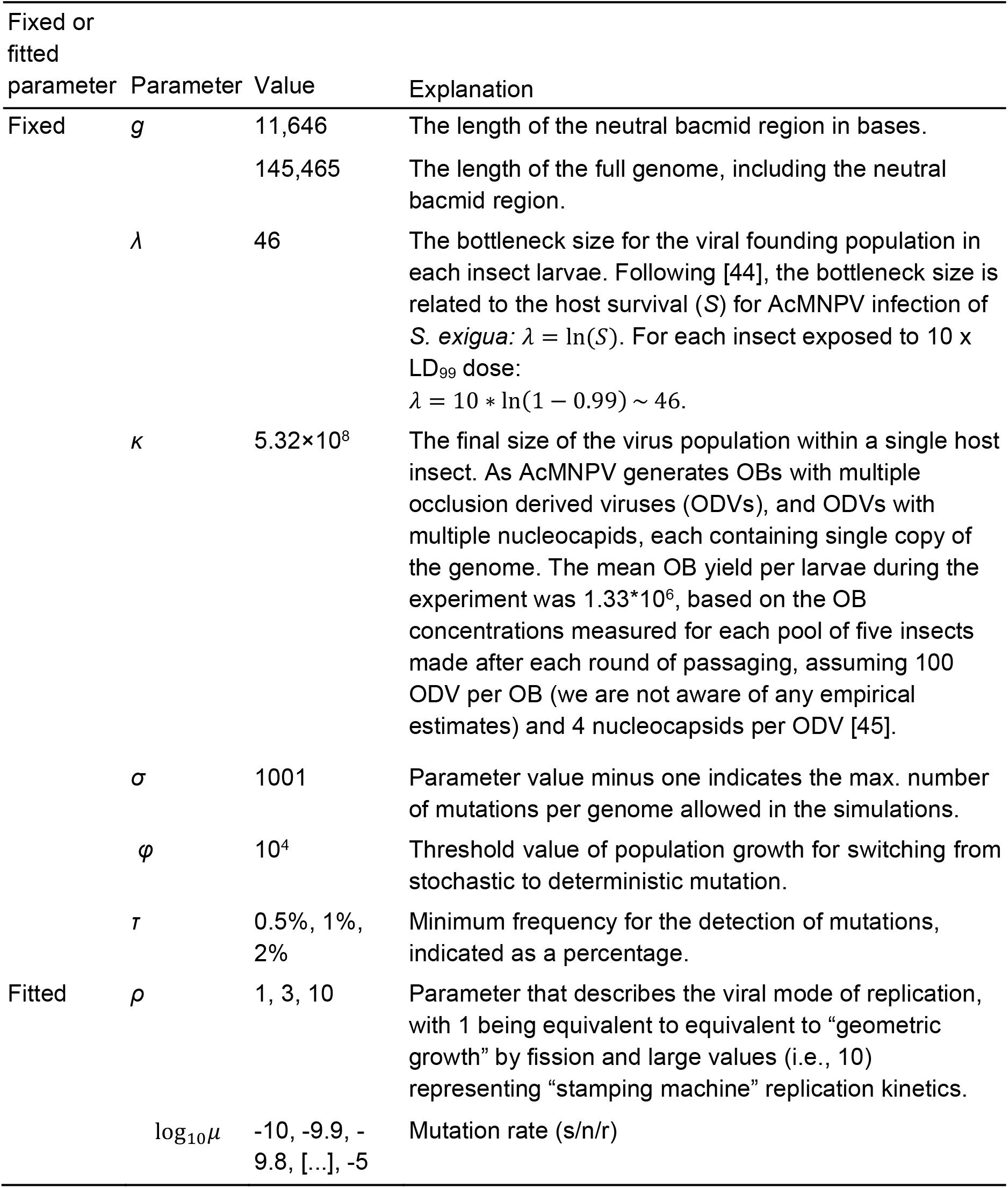
Parameters for the models for estimating mutation rates

At the start of the infection of an individual host, there is a bottleneck with *λ* virus genomes initiating infection. For each position in the genome, we draw the number of genomes containing a mutation at this position from a binomial distribution, where for the *i*^th^ position: 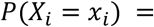 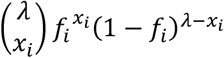, where *X* is the number of mutant genomes added to the population at a particular step in the infection process (and *f* is the frequency of mutated bases at position *i*). We obtained an estimate of *λ* = 46 by considering the relationship between host mortality and the number of viral founders [44] (see Table 3). The virus population then expands within the host exponentially with a replication factor *ρ* per cycle of virus replication within the host, such that *N*_*t*_ = *λ*(1 + *ρ*)^*t*^ where *N* is the number of genomes present at a time *t*, measured in generations of viral replication within the host. It is unknown what the mode of replication [30,31] is for a baculovirus. We therefore used values of 1 (“geometric” or “symmetric” replication with a doubling of the number of copies per cycle – one original genome copy and one replicated copy), 3 (“mixed” replication – one original genome copy and three replicated copies) and 10 (“stamping machine” or “asymmetric” replication) for *ρ*. Replication proceeds until the carrying capacity *κ* of a host is reached, with an expansion to exactly *κ* virus genomes allowed in the final round of replication. During each round of replication, the number of new mutants that occur at each position follows a binomial distribution, such that the mutation rate *μ* is the probability of success and *η*, the number of genomes generated during that round of replication which are not mutated at this nucleotide position (i.e., *η*_*i,t*_ = *ρN*_*t*-1_(1 - *f*_*i,t*-1_)), is the number of events: 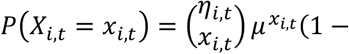 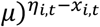. However, to make the model computationally tractable for large population sizes, we switched to deterministic mutation (I.e., *X*_*i,t*_ = *η*_*i,t*_*μ*) once a large number of virus genomes was being generated relative to the mutation rate (*N*_*t*_*φ* > *μ*^−1^), where *φ* is a constant. Note that all mutations are assumed to be neutral and that the bottleneck at the start of infection is narrow (*λ* = 46). Mutations occurring late in infection when the viral census population size is large therefore will rarely be sampled during the bottleneck events at the start of the next round of infection (i.e. serial transfer). To model the infection of five host larvae per replicate, we simulated five separate infections and pooled the viral progeny by taking the mean mutation frequency per site over the five larvae for each position in the genome, and using these *fi* values for the next round of infection.

To fit the model to the data, we first ran 1000 simulations for each combination of parameter values (i.e., *μ*, *ρ* and *τ*) to generate model predictions. We compared the observed number of experimental replicates (*q*_*j*_) with *j* – *1* mutated nucleotide positions with a frequency *f*_*i*_ higher than the threshold value *τ*, to the frequency predicted by the model (*β*_*j*_). (I.e., *q*_1_ is the number of experimental replicates with 0 nucleotide positions for which *f*_*i*_ > *τ*, *q*_2_ is the number of replicates with 1 nucleotide positions for which *f*_*i*_ > *τ,* etc.) The multinomial pseudo-likelihood of any realization is then: 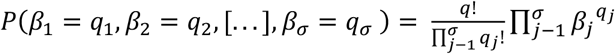, where *σ-1* = 1000 is the maximum number of mutated bases that were tracked in the simulations. If any values of *β* exceeded *σ*-1 in any simulation, results from that set of parameter values were excluded from further analysis. We fitted the model to 1000 bootstrapped datasets to determine the 95% confidence interval of the parameter estimates.

### Mutation rate estimates with established approaches

We estimated the mutation rate with an established approach, to compare to the estimates made with our simulation-model approach. A canonical approach [9] for calculating mutation rate (s/n/r) is 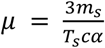, where *m*_*s*_ is the number of observed mutations observed in sequenced clones, *T*_*s*_ is the mutational target size, *c* is the number of viral generations, and *α* is a correction for the effects of selection. To obtain *m*_*s*_ from our deep sequencing data, we sum the frequency of all observed mutations (*f*_*i*_) above the threshold for mutation selection *τ* in all lineages. If these sequencing data are accurate and mutations are neutral, the frequency of each mutation is also the probability that this mutation would be detected in a randomly selected clone by sequencing. This approach is a simple approximation, as here we do not consider the effect of the threshold for mutation detection (*τ*) on mutation rate estimates. (Lowering *τ* will lead to a larger number of mutations and consequently a higher mutation rate estimate. To keep this method as simple as possible and free of additional assumptions, we choose not to incorporate any corrections.) Note that because we sum the frequency of all possible mutations over each site, we can drop the three in the numerator. To obtain *T*_*s*_, we multiply the length of the neutral region by the number of replicates. To obtain *c* we estimate the number of generations assuming different values for *ρ* (i.e., 1, 3 and 10 as for the simulation-based model fitting), such that *c* = *θ* · *ln*(*κ*/*λ*)/*ln*(1 + *ρ*), where *θ* is the number of passages. Finally, we can drop *α* because we only consider mutations in the neutral bacmid region. One-thousand bootstrapped datasets were used to obtain fiducial limits for the mutation rate estimates.

### dI/dS and dN/dS analyses

Estimates of dN/dS (i.e., the normalized rate of nonsynonymous mutations, here made for authentic viral genes) were made using standard methods [46,47]. As we are not aware of any estimates of the mutation spectrum for insect DNA viruses and our own data suggest these biases may not be very strong (Table 2), we assume no mutation bias is present (i.e., a transition to transversion ratio of 1). For the dI/dS [3], the dS term is the same as for the dN/dS analysis, derived from the results for authentic viral genes. The dI term is determined for mutations in the bacmid neutral region. Ninety-five percent confidence intervals of the dI/dS and dN/dS were obtained using 1000 bootstrapped datasets, and data were tested for significance with a one-sample *t*-test on the dN/dS values calculated for individual samples compared to a test value of 1. However, when *τ* > 0.5% for one or more samples the number of synonymous samples was 0, and hence these analyses could only be performed for *τ* = 0.5%. Full results and R code are available in the Supplementary Online Information (Appendices 7-9).

## Acknowledgements

The authors thank Yue Han and Marleen Henkens for technical assistance. G.A. was supported by the Netherlands Fellowship Program PhD Grant No. CF7554/2011 (Nuffic, www.nuffic.nl). MPZ was supported by a VIDI grant from the Netherlands Organization for Scientific Research Grant No. 016.VIDI.171.061 (NWO, www.nwo.nl).

The funders had no role in study design, data collection and analysis, decision to publish, or preparation of the manuscript. The authors have declared that no competing interests exist.

## Appendix

**Table S1:** Relative frequencies of mutations observed per type, for the whole genome and bacmid only datasets, across mutation frequency threshold values (*τ*), given as a percentage.

| Mutation | Relative frequency of mutation |  |  |  | Mutation type |
| --- | --- | --- | --- | --- | --- |
|  | Whole genome |  |  | Bacmid only |  |
| | $\tau = 0.5$<br>(n=69) | $\tau = 1$<br>(n=20) | $\tau = 2$<br>(n=8) | $\tau = 0.5$<br>(n=9) | |
| CA | 0.174 | 0.250 | 0.125 | 0.111 | Transversion |
| CG | 0.014 | 0.000 | 0.000 | 0.000 | Transversion |
| TA | 0.043 | 0.000 | 0.000 | 0.000 | Transversion |
| TG | 0.014 | 0.000 | 0.000 | 0.111 | Transversion |
| AC | 0.043 | 0.000 | 0.000 | 0.111 | Transversion |
| AT | 0.058 | 0.000 | 0.000 | 0.111 | Transversion |
| GC | 0.029 | 0.000 | 0.000 | 0.000 | Transversion |
| GT | 0.217 | 0.300 | 0.125 | 0.111 | Transversion |
| GA | 0.130 | 0.200 | 0.250 | 0.222 | Transition |
| CT | 0.101 | 0.150 | 0.250 | 0.111 | Transition |
| TC | 0.072 | 0.050 | 0.125 | 0.000 | Transition |
| AG | 0.101 | 0.050 | 0.125 | 0.111 | Transition |

**Figure S1:**
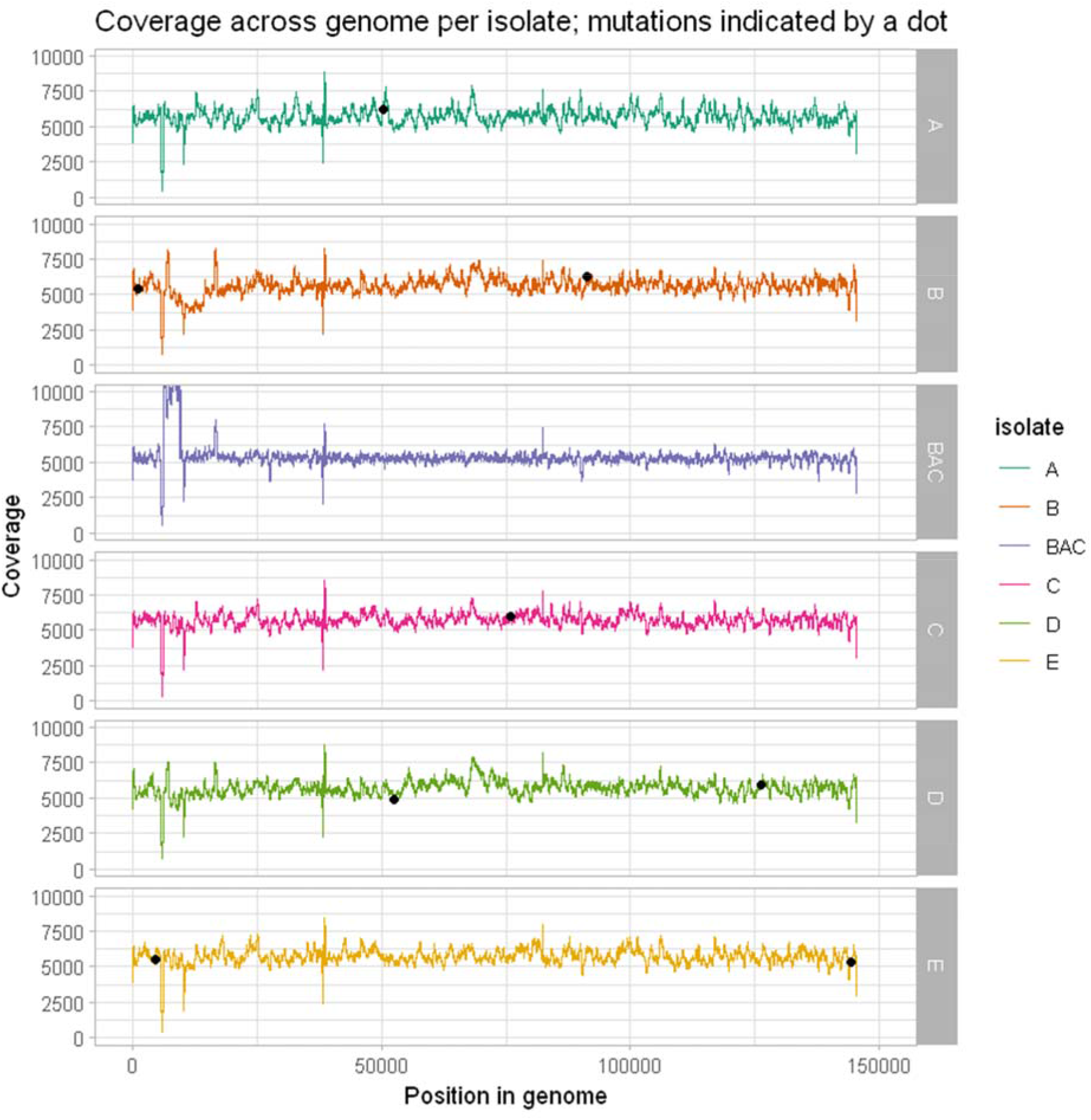
We show coverage along the genome for each evolved line (A, B, C, D and E) as well as ancestral strain BAC. Position of mutations observed at mutation frequency threshold value (*τ*) = 2 are shown as black dots. Coverage patterns are similar between the different isolates. The peak observed at around 10000 bp for the BAC isolate is due to the presence of empty bacmid vectors in sequencing data and is omitted from mutation calling.

**Figure S2:**
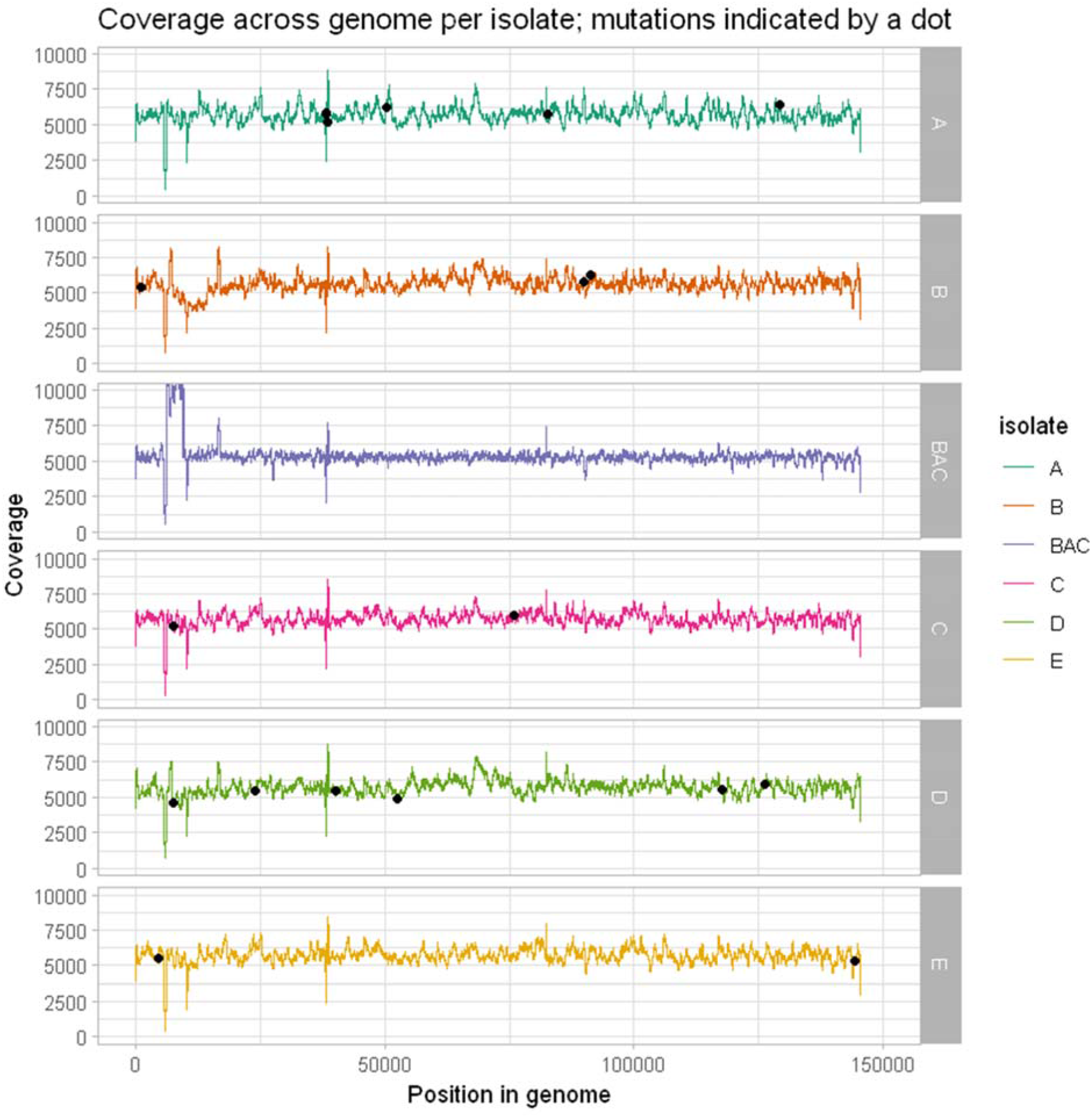
We show coverage along the genome for each evolved line (A, B, C, D and E) as well as ancestral strain BAC. Position of mutations observed at mutation frequency threshold value (*τ*) = 1 are shown as black dots. Coverage patterns are similar between the different isolates. The peak observed at around 10000 bp for the BAC isolate is due to the presence of empty bacmid vectors in sequencing data and is omitted from mutation calling.

**Figure S3:**
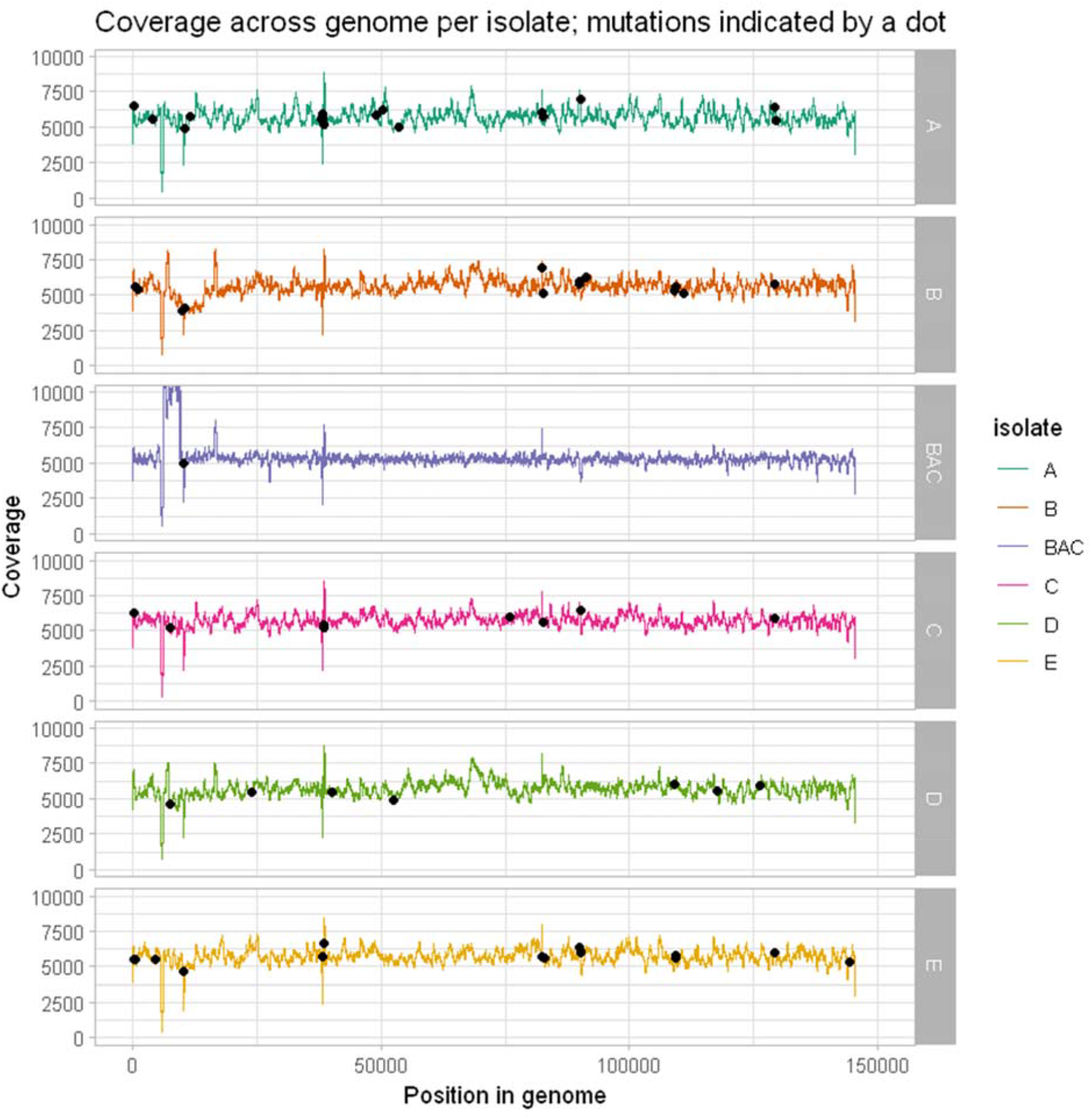
We show coverage along the genome for each evolved line (A, B, C, D and E) as well as ancestral strain BAC. Position of mutations observed at mutation frequency threshold value (*τ*) = 0.5 are shown as black dots. Coverage patterns are similar between the different isolates. The peak observed at around 10000 bp for the BAC isolate is due to the presence of empty bacmid vectors in sequencing data and is omitted from mutation calling.

**Figure S4:**
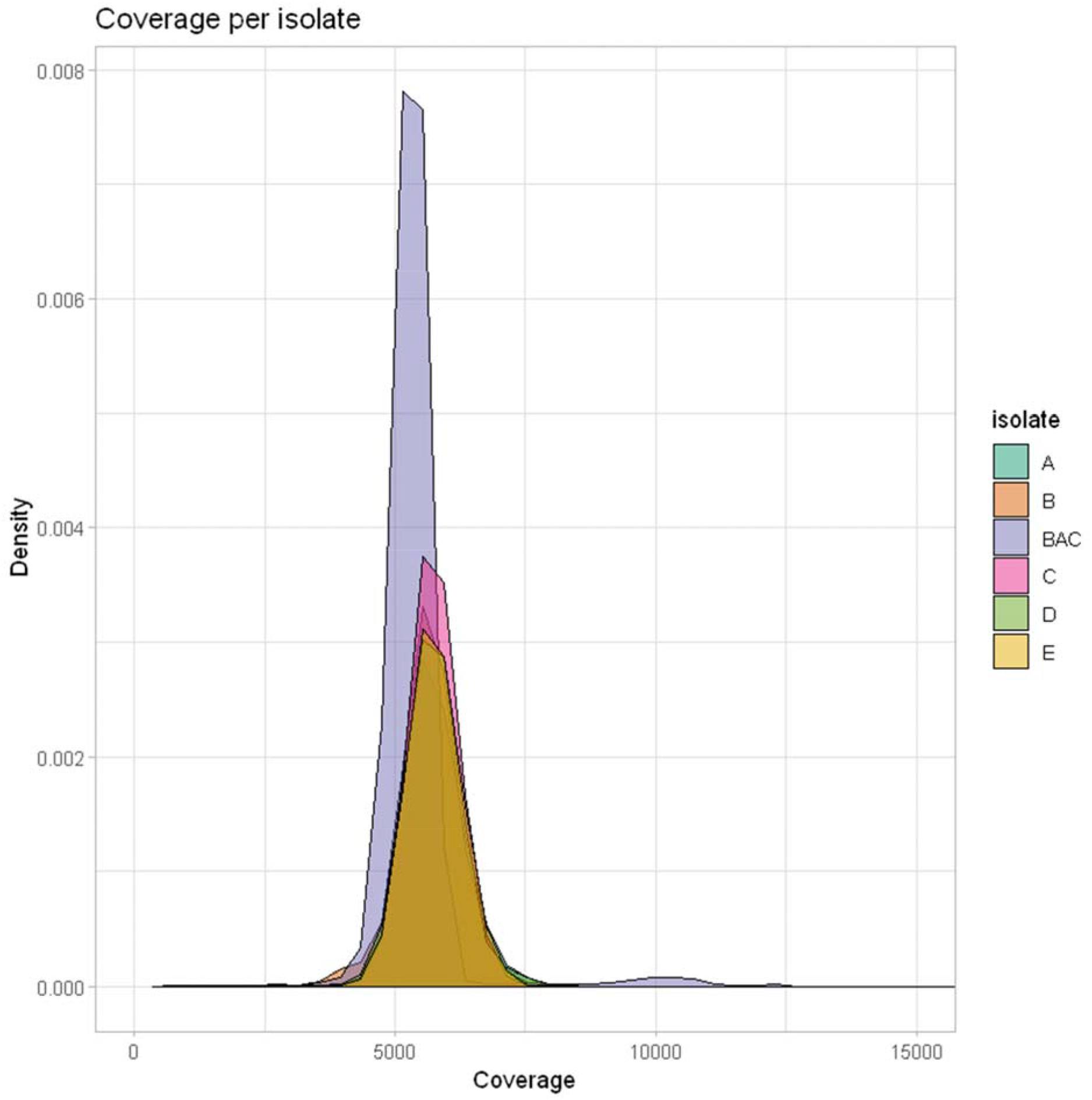
Coverage distribution per isolate after subsampling to approximately equal mean coverage. Isolates have a mean coverage of around 5500. The BAC isolate is showing an additional peak at a coverage of around 10000, which is explained by the presence of empty bacmid vectors in sequencing data.

**Figure S5:**
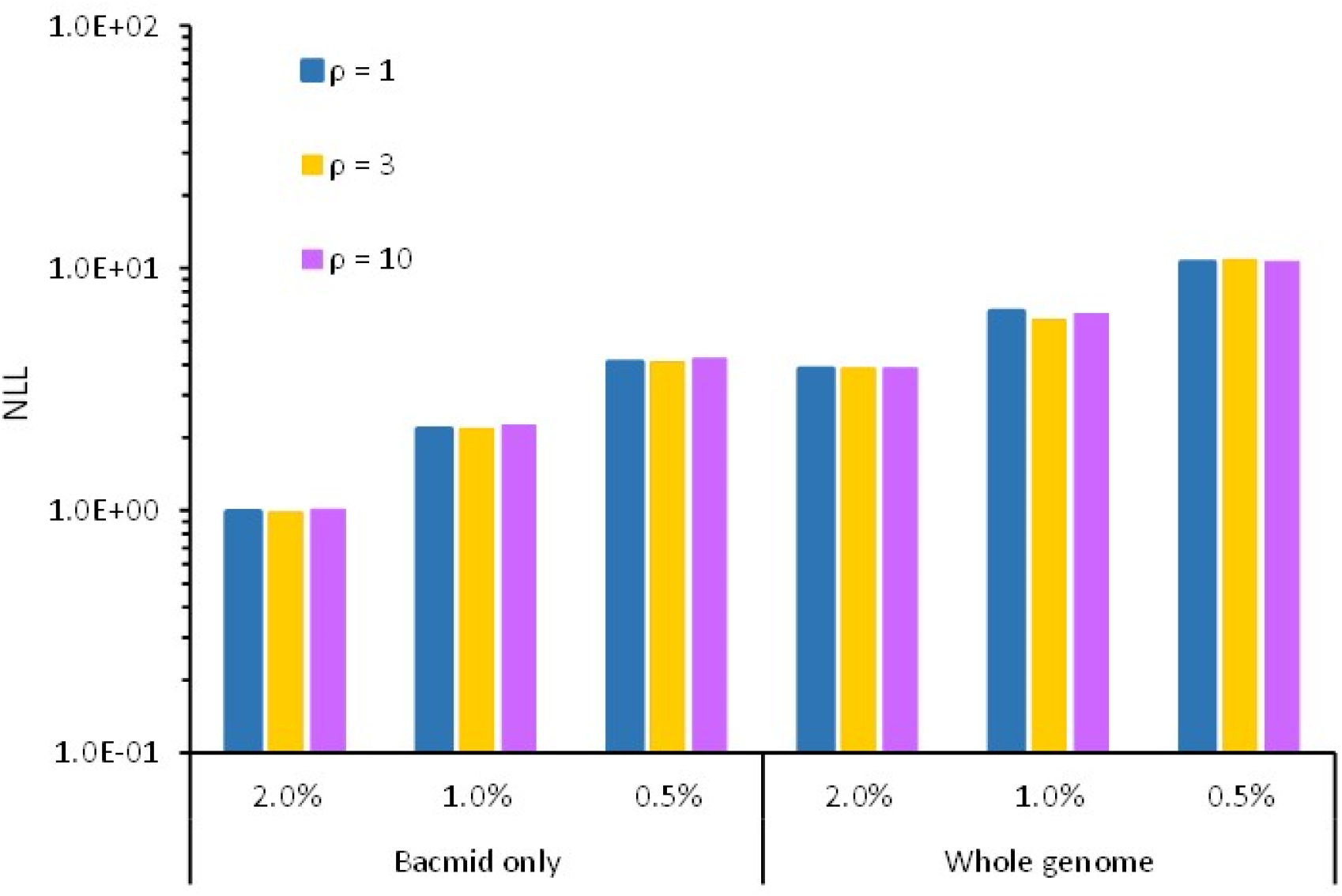
We show the negative log likelihood (NLL) for models fitted with different mutation frequency threshold values (*τ*), given as a percentage. Mutation frequency threshold values clearly effect model fit, as they have an effect on the number of mutations that will be detected. By contrast, assumptions on the value for the parameter that determines the mode of virus replication (*ρ*) had little effect on model fit. This result is not surprising however, given that our model does not consider the frequency of mutations, but simply the number of bases with a mutation frequency greater than *τ.*

**Figure S6:**
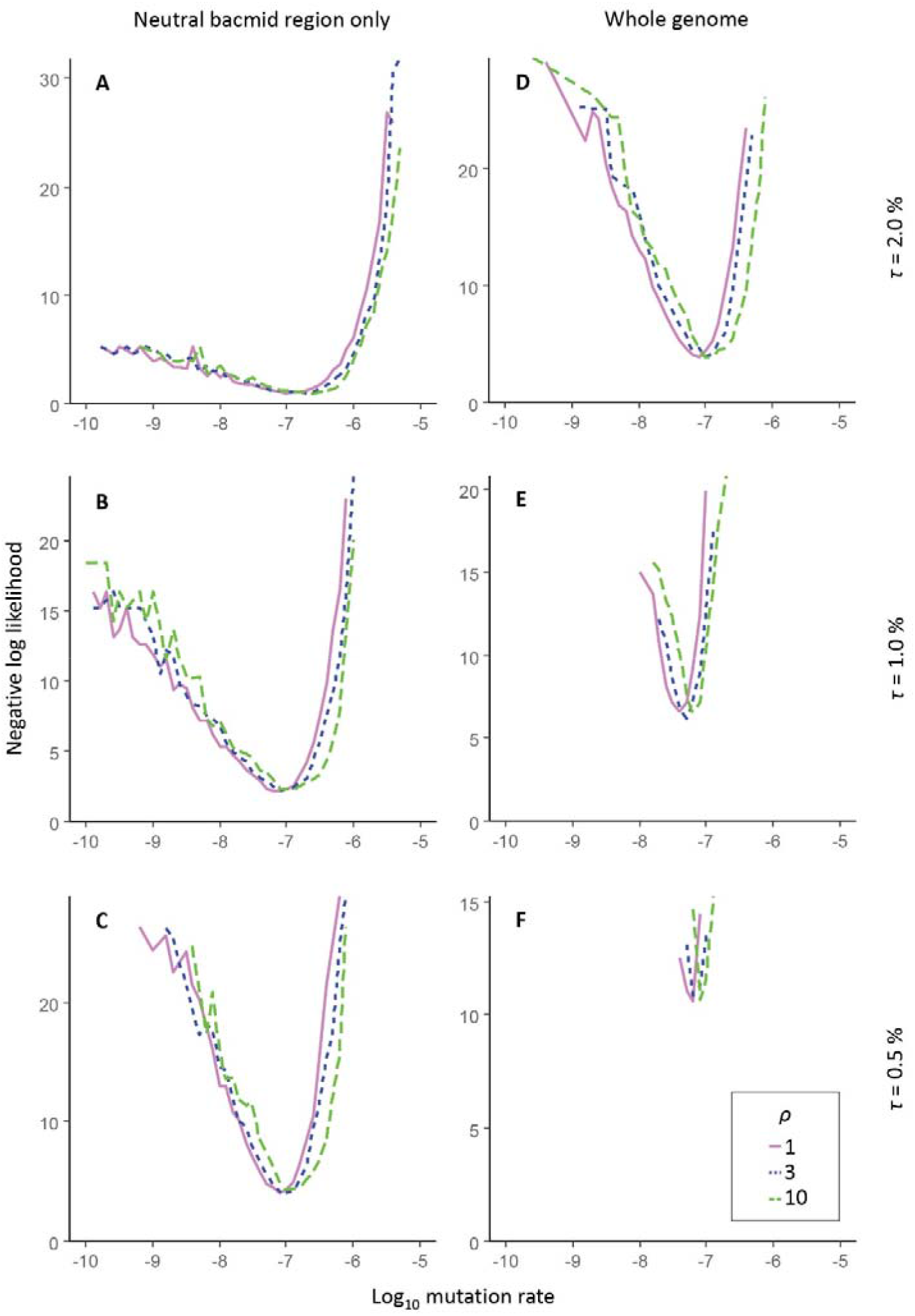
Results of the fitting of the simulation model to the experimental data, for making mutation rate estimates. For all panels, the abscissa is the log10 mutation rate and the originate the negative log likelihood. Panels on the left correspond to the neutral bacmid region data, and panels on the right to the whole genome data. Different thresholds for the detection of mutations (*τ*) were also employed, ranging from 0.5% (bottom row) to 1% (middle row) to 2% (top row). The different colors and line types indicate model fit for different values of *ρ*, the parameter that describes the viral mode of replication. Whereas higher values of *ρ* lead to lower mutation rate estimates, this model parameter does not impact the minimum for the NLL.

**Figure S7:**
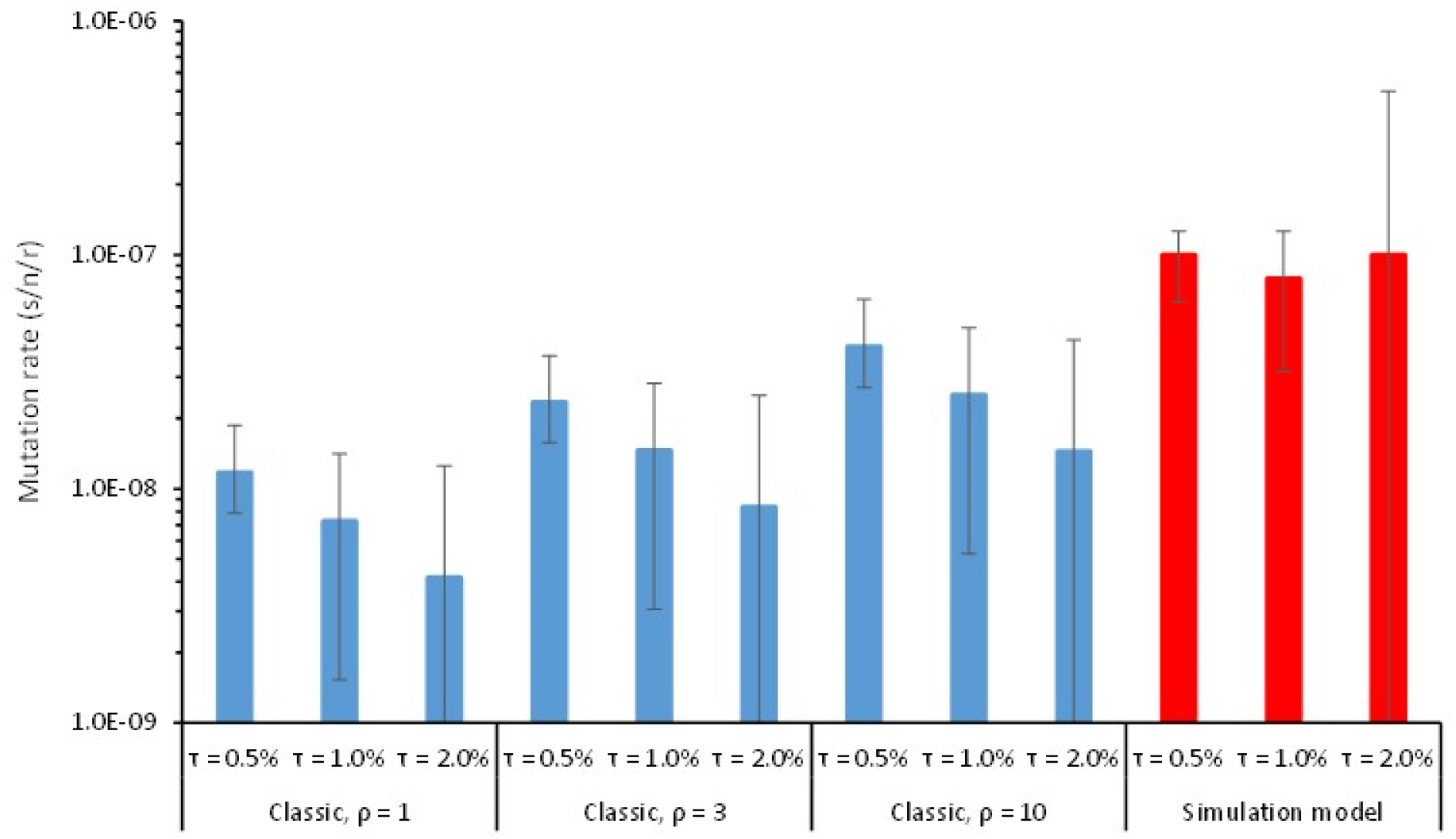
Estimate of mutation rate (s/n/r) using established methods applied to deep sequencing data (blue bars, samples categorized as “Classic”), as an alternative to our approach using a simulation-based model (red bars, samples categorized as “Simulation model”). Mutation rates were estimated for different values of the viral mode of replication (*ρ*) and with different values for the threshold of mutation detection (*τ*). Error bars represent the 95% fiducial limits, as determined by bootstrapping. When the lower fiducial limit extends beyond the lower limit of the axis, this indicates a lower fiducial limit of zero. Overall, these estimates were lower than those obtained with the approach employing a simulation model. As baculoviruses most likely employ rolling circle amplification, replication is likely to have a high value of *ρ*. Furthermore, this method is clearly sensitive to differences in the mutation detection threshold, as the cumulative frequency of mutations above this value is used required to estimate mutations and no correction is made for this threshold. Therefore, the best estimates with this approach assume the highest value of *ρ* and the lowest mutation detection thresholds (*τ*), provided all mutations are assumed to be bona fide. The estimate for these conditions (*τ* = 0.5% and *ρ* = 10), renders an estimate of *μ* = 4 × 10^−8^ which is lower but roughly similar than for our simulation-based approach (*μ* ~ 10^−7^).

## Notes

### Competing Interest Statement

The authors have declared no competing interest.

### Summary of Updates

The complete supplementary materials were added to this version, as these were not available in the version cross-posted by the journal.

## References

1. de Visser JAGM, Krug J. Empirical fitness landscapes and the predictability of evolution. Nat Rev Genet. 2014;15: 480–490.

2. Bull JJ, Badgett MR, Wichman HA, Huelsenbeck JP, Hillis DM, Gulati A, et al. Exceptional convergent evolution in a virus. Genetics. 1997;147: 1497–1507.

3. Tenaillon O, Rodriguez-Verdugo A, Gaut RL, McDonald P, Bennett AF, Long AD, et al. The molecular diversity of adaptive convergence. Science. 2012;335: 457–461.

4. Sprouffske K, Aguílar-Rodríguez J, Sniegowski P, Wagner A. High mutation rates limit evolutionary adaptation in *Escherichia coli*. PLOS Genet. 2018;14: 1–31.

5. Van Dijk T, Hwang S, Krug J, De Visser JAGM, Zwart MP. Mutation supply and the repeatability of selection for antibiotic resistance. Phys Biol. 2017;14: 055005.

6. Salverda MLM, Koomen J, Koopmanschap B, Zwart MP, De Visser JAGM. Adaptive benefits from small mutation supplies in an antibiotic resistance enzyme. Proc Natl Acad Sci U S A. 2017;114: 12773–12778.

7. Peck KM, Lauring AS. Complexities of viral mutation rates. J Virol. 2018;92: 1–8.

8. Hughes D, Andersson DI. Evolutionary trajectories to antibiotic resistance. Annu Rev Microbiol. 2017;71: 579–596.

9. Sanjuán R, Nebot MR, Chirico N, Mansky LM, Belshaw R. Viral mutation rates. J Virol. 2010;84: 9733–9748.

10. Sanjuán R, Domingo-Calap P. Mechanisms of viral mutation. Cell Mol Life Sci. 2016;73: 4433–4448.

11. Luria SE. The frequency distribution of spontaneous bacteriophage mutants as evidence for the exponential rate of phage reproduction. Cold Spring Harb Symp Quant Biol. 1951;16: 463–470.

12. Pauly MD, Procario MC, Lauring AS. A novel twelve class fluctuation test reveals higher than expected mutation rates for Influenza A viruses. Elife. 2017;6: 1–18.

13. Sanjuan R, Moya A, Elena S. The distribution of fitness effects caused by single-nucleotide substitutions in an RNA virus. Proc Natl Acad Sci USA. 2004;101: 8396–8401.

14. Carrasco P, de la Iglesia F, Elena SF. Distribution of fitness and virulence effects caused by single-nucleotide substitutions in Tobacco etch virus. J Virol. 2007;81: 12979–12984.

15. Mahmoudabadi G, Phillips R. A comprehensive and quantitative exploration of thousands of viral genomes. Elife. 2018;7: 1–26.

16. Knopf CW. Evolution of viral DNA-dependent DNA polymerases. Virus Genes. 1998;16: 47–58.

17. Gago S, Elena SF, Flores R, Sanjuán R. Extremely high mutation rate of a hammerhead viroid. Science. 2009;323: 1308.

18. Haase S, Sciocco-Cap A, Romanowski V. Baculovirus insecticides in Latin America: Historical overview, current status and future perspectives. Viruses. 2015;7: 2230–2267.

19. van Oers MM. Opportunities and challenges for the baculovirus expression system. J Invertebr Pathol. 2011;107: S3–S15.

20. Harrison R, Herniou E, Jehle J, Theilmann D, Burand J, Becnel J, et al. ICTV virus taxonomy profile: Baculoviruses. J Gen Virol. 2018;99: 1185–1186.

21. Smith IR, Crook NE. In vivo isolation of baculovirus genotypes. Virology. 1988;166: 240–244.

22. Cory JS, Green BM, Paul RK, Hunter-Fujita F. Genotypic and phenotypic diversity of a baculovirus population within an individual insect host. J Invertebr Pathol. 2005;89: 101–111.

23. Kennedy DA, Dwyer G. Effects of multiple sources of genetic drift on pathogen variation within hosts. PLoS Biol. 2018;16: e2004444.

24. Chateigner A, Bézier A, Labrousse C, Jiolle D, Barbe V, Herniou EA. Ultra deep sequencing of a baculovirus population reveals widespread genomic variations. Viruses. 2015;7: 3625–3646.

25. Sanjuán R, Agudelo-Romero P, Elena SF. Upper-limit mutation rate estimation for a plant RNA virus. Biol Lett. 2009;5: 394–396.

26. Tromas N, Elena SF. The rate and spectrum of spontaneous mutations in a plant RNA virus. Genetics. 2010;185: 983–989.

27. Luckow VA, Lee SC, Barry GF, Olins PO. Efficient generation of infectious recombinant baculoviruses by site-specific transposon-mediated insertion of foreign genes into a baculovirus genome propagated in *Escherichia coli*. J Virol. 1993;67: 4566–4579.

28. Willemsen A, Zwart MP. On the stability of sequences inserted into viral genomes. Virus Evol. 2019;5: 1–16.

29. Sardanyés J, Solé R V., Elena SF. Replication mode and landscape topology differentially affect RNA virus mutational load and robustness. J Virol. 2009;83: 12579–12589.

30. Martínez F, Sardanyés J, Elena SF, Daròs JA. Dynamics of a plant RNA virus intracellular accumulation: Stamping machine vs. geometric replication. Genetics. 2011;188: 637–646.

31. Schulte MB, Draghi JA, Plotkin JB, Andino R. Experimentally guided models reveal replication principles that shape the mutation distribution of RNA viruses. Elife. 2015;4: 1–18.

32. Leisy DJ, Rohrmann GF. Characterization of the replication of plasmids containing *hr* sequences in baculovirus-infected *Spodoptera frugiperda* cells. Virology. 1993;196: 722–730.

33. Sanjuan R. Viral mutation rate estimates. [cited 26 Feb 2021]. Available: http://www.uv.es/rsanjuan/virmut

34. Blanco L, Bernad A, Blasco MA, Salas M. A general structure for DNA-dependent DNA polymerases. Gene. 1991;100: 27–38.

35. Tomalski MD, Wu J, Miller LK. The location, sequence, transcription, and regulation of a baculovirus DNA polymerase gene. Virology. 1988;167: 591–600.

36. Hang X, Guarino LA. Purification of Autographa californica nucleopolyhedrovirus DNA polymerase from infected insect cells. J Gen Virol. 1999;80: 2519–2526.

37. Karam JD, Konigsberg WH. DNA polymerase of the T4-related bacteriophages. Prog Nucleic Acid Res Mol Biol. 2000;64: 65–96.

38. Acevedo A, Brodsky L, Andino R. Mutational and fitness landscapes of an RNA virus revealed through population sequencing. Nature. 2014;505: 686–690.

39. Simon O, Williams T, Caballero P, Lopez-Ferber M. Dynamics of deletion genotypes in an experimental insect virus population. Proc R Soc B: Biol Sci. 2006;273: 783–790.

40. Clavijo G, Williams T, Simon O, Munoz D, Cerutti M, Lopez-Ferber M, Caballero P. Mixtures of complete and *pif1*- and *pif2*-deficient genotypes are required for increased potency of an insect nucleopolyhedrovirus. J Virol. 2009;83: 5127–5136.

41. Anonymous. Seqtk: Toolkit for processing sequences in FASTA/Q formats. [cited July 7 2021] Available: https://github.com/lh3/seqtk.

42. Anonymous. CLC Genomics Workbench 20.0. [cited 8 Apr 2021]. Available: https://digitalinsights.qiagen.com

43. The R Foundation for Statistical Computing. R: A language and environment for statistical computing. Vienna, Austria; 2020.

44. Zwart MP, Hemerik L, Cory JS, de Visser JAGM, Bianchi FJJA, van Oers MM, Vlak JM, Hoekstra RF, van der Werf W. An experimental test of the independent action hypothesis in virus-insect pathosystems. Proc R Soc B: Biol Sci. 2009;276: 2233–42.

45. Zwart MP, van Oers MM, Cory JS, van Lent JWM, van der Werf W, Vlak JM. Development of a quantitative real-time PCR for determination of genotype frequencies for studies in baculovirus population biology. J Virol Methods. 2008;148: 146–54.

46. Yang Z, Bielawski JR. Statistical methods for detecting molecular adaptation. Trends Ecol Evol. 2000;15: 496–503.

47. Zwart MP, Ali G, van Strien EA, Schijlen EGWM, Wang M, van der Werf W, et al. Identification of loci associated with enhanced virulence in Spodoptera litura nucleopolyhedrovirus isolates using deep sequencing. Viruses. 2019;11: 1–13.

