## Supplementary material for "Empirical estimates of the mutation rate for an alphabaculovirus": Appendix 1 CLC GW Settings.pdf

### Workflow_10_2020

| Workflow Input (FastBacDual_polh_1805_annotated) | |
|---|---|
| Workflow Input | FastBacDual_polh_1805_annotated |
| Import Command | |

| Map Reads to Reference | |
|---|---|
| References | Defined by: Workflow Input ( FastBacDual_polh_1805_annotated) |
| Masking mode | No masking |
| Masking track | |
| Match score | 1 |
| Mismatch cost | 2 |
| Cost of insertions and deletions | Linear gap cost |
| Insertion cost | 3 |
| Deletion cost | 3 |
| Insertion open cost | 6 |
| Insertion extend cost | 1 |
| Deletion open cost | 6 |
| Deletion extend cost | 1 |
| Length fraction | 0.5 |
| Similarity fraction | 0.8 |
| Global alignment | false |
| Auto-detect paired distances | true |
| Non-specific match handling | Map randomly |

| Low Frequency Variant Detection | |
|---|---|
| Required significance (%) | 1.0 |
| Ignore positions with coverage above | 100000 |
| Restrict calling to target regions | |
| Ignore broken pairs | true |
| Ignore non-specific matches | Reads |
| Minimum read length | 20 |
| Minimum coverage | 10 |
| Minimum count | 2 |
| Minimum frequency (%) | 0.5 |

| Low Frequency Variant Detection | |
|---|---|
| Base quality filter | true |
| Neighborhood radius | 5 |
| Minimum central quality | 20 |
| Minimum neighborhood quality | 15 |
| Read direction filter | false |
| Direction frequency (%) | 5.0 |
| Relative read direction filter | true |
| Significance (%) | 1.0 |
| Read position filter | false |
| Significance (%) | 1.0 |
| Remove pyro-error variants | false |
| In homopolymer regions with minimum length | 3 |
| With frequency below | 0.8 |

| Trim Reads | |
|---|---|
| Quality trim | true |
| Quality limit | 0.05 |
| Ambiguous trim | true |
| Ambiguous limit | 2 |
| Trim adapter list | |
| Automatic read-through adapter trimming | true |
| Trim homopolymers from 5' | false |
| Trim homopolymers from 3' | false |
| polyA | false |
| polyC | false |
| polyG | true |
| polyT | false |
| Remove 5' terminal nucleotides | false |
| Number of 5' terminal nucleotides | 1 |
| Remove 3' terminal nucleotides | false |
| Number of 3' terminal nucleotides | 1 |
| Fixed length trimming | false |
| Maximum length | 150 |
| Trim from side | 3'-end |

| Trim Reads | |
|---|---|
| Discard short reads | true |
| Minimum number of nucleotides in reads | 15 |
| Discard long reads | true |
| Maximum number of nucleotides in reads | 1000 |

| QC for Read Mapping | |
|---|---|
| Long contigs threshold | 10000 |
| Short contigs threshold | 200 |
