## Supplementary material for "Empirical estimates of the mutation rate for an alphabaculovirus": Appendix 2 Subsampling MutationCalling v3 freq=0.5.html


### AcMNPV mutation rate estimation (with subsampling to equal coverage)¶

Authors: Dieke Boezen & Mark Zwart
Date: September and October 2020

This project aims to determine the mutationrate for baculovirus AcMNPV using Illumina NGS data from ancestral (BAC) and evolved lines (A-E) of AcMNPV in *Spodoptera exigua/litura*. The pipeline in this notebook filters mutations from variant calling output of CLC genomics workbench 20, and makes plots of the coverage across the genome and the mutations present. Corresponding to this pipeline for the "empirical" component, there is a mutation rate model fitted to the mutations called here.

### 0. Subsampling NGS data with seqtk to equal coverage (unix on command line)¶

**seqtk sample** allows us to subsample reads from a fastq file. Set seed to 42 to ensure read pairs are not lost when subsampling, and sample 2.8M reads per file to ge approx 5.6M reads mapped to the reference genome, which will be comparable to the mapped reads for the BAC sample. Subsampled files were rerun in the CLC pipeline.

In [74]:

```
(subsamplingbaculo) diekeb@nioo0025:~/subsampled_reads_AcMNPV$ seqtk sample -s42 /mnt/nfs/bioinfdata/ngs/ME/zwart_group/RawData/IlluminaNGS/AcMNPV_evolution/A_H75TJDMXX_L1_1.clean.fq 2800000 > sub_A_1.fq
(subsamplingbaculo) diekeb@nioo0025:~/subsampled_reads_AcMNPV$ seqtk sample -s42 /mnt/nfs/bioinfdata/ngs/ME/zwart_group/RawData/IlluminaNGS/AcMNPV_evolution/A_H75TJDMXX_L1_2.clean.fq 2800000 > sub_A_2.fq
(subsamplingbaculo) diekeb@nioo0025:~/subsampled_reads_AcMNPV$ seqtk sample -s42 /mnt/nfs/bioinfdata/ngs/ME/zwart_group/RawData/IlluminaNGS/AcMNPV_evolution/B_H77KHDMXX_L1_1.clean.fq 2800000 > sub_B_1.fq
(subsamplingbaculo) diekeb@nioo0025:~/subsampled_reads_AcMNPV$ seqtk sample -s42 /mnt/nfs/bioinfdata/ngs/ME/zwart_group/RawData/IlluminaNGS/AcMNPV_evolution/B_H77KHDMXX_L1_2.clean.fq 2800000 > sub_B_2.fq
(subsamplingbaculo) diekeb@nioo0025:~/subsampled_reads_AcMNPV$ seqtk sample -s42 /mnt/nfs/bioinfdata/ngs/ME/zwart_group/RawData/IlluminaNGS/AcMNPV_evolution/C_H75TJDMXX_L1_1.clean.fq 2800000 > sub_C_1.fq
(subsamplingbaculo) diekeb@nioo0025:~/subsampled_reads_AcMNPV$ seqtk sample -s42 /mnt/nfs/bioinfdata/ngs/ME/zwart_group/RawData/IlluminaNGS/AcMNPV_evolution/C_H75TJDMXX_L1_2.clean.fq 2800000 > sub_C_2.fq
(subsamplingbaculo) diekeb@nioo0025:~/subsampled_reads_AcMNPV$ seqtk sample -s42 /mnt/nfs/bioinfdata/ngs/ME/zwart_group/RawData/IlluminaNGS/AcMNPV_evolution/D_H77KHDMXX_L1_1.clean.fq 2800000 > sub_D_1.fq
(subsamplingbaculo) diekeb@nioo0025:~/subsampled_reads_AcMNPV$ seqtk sample -s42 /mnt/nfs/bioinfdata/ngs/ME/zwart_group/RawData/IlluminaNGS/AcMNPV_evolution/D_H77KHDMXX_L1_2.clean.fq 2800000 > sub_D_2.fq
(subsamplingbaculo) diekeb@nioo0025:~/subsampled_reads_AcMNPV$ seqtk sample -s42 /mnt/nfs/bioinfdata/ngs/ME/zwart_group/RawData/IlluminaNGS/AcMNPV_evolution/E_H7FJMDMXX_L1_1.clean.fq 2800000 > sub_E_1.fq
(subsamplingbaculo) diekeb@nioo0025:~/subsampled_reads_AcMNPV$ seqtk sample -s42 /mnt/nfs/bioinfdata/ngs/ME/zwart_group/RawData/IlluminaNGS/AcMNPV_evolution/E_H7FJMDMXX_L1_2.clean.fq 2800000 > sub_E_2.fq
```

```
Error in parse(text = x, srcfile = src): <text>:1:21: unexpected symbol
1: (subsamplingbaculo) diekeb
                        ^
Traceback:
```

### 1. Load libraries¶

In [2]:

```
library(tidyverse)
library(janitor)
library(RColorBrewer)
```

### 2. Coverage¶

In the section below, I make some plots of the coverage across the reference genome per each of the strains/isolates to check for odd patterns and get an idea of variation within and between isolates.

#### Load coverage data and clean the dataset¶

Data is loaded as seperate TSV files containing the coverage per position in the reference genome as generated in CLC genomics wb 20. The dataset is mutated to include the isolate information, and merged to generate plots. We also generate ranks for the coverage so we can exclude "extremes" in the downstream variant calling.

In [3]:

```
df_BAC_cov <- read_tsv("C:/Users/Dieke/Google Drive/NIOO PhD/AcMNPV_mutationrate/coverage/BAC_H752KDMXX_L1_1.clean (paired, trimmed pairs) mapping.tsv", col_names = FALSE, col_types = cols())
df_A_cov <- read_tsv("C:/Users/Dieke/Google Drive/NIOO PhD/AcMNPV_mutationrate/coverage/sub_A_1 (paired, trimmed pairs) mapping.tsv", col_names = FALSE, col_types = cols())
df_B_cov <- read_tsv("C:/Users/Dieke/Google Drive/NIOO PhD/AcMNPV_mutationrate/coverage/sub_B_1 (paired, trimmed pairs) mapping.tsv", col_names = FALSE, col_types = cols())
df_C_cov <- read_tsv("C:/Users/Dieke/Google Drive/NIOO PhD/AcMNPV_mutationrate/coverage/sub_C_1 (paired, trimmed pairs) mapping.tsv", col_names = FALSE, col_types = cols())
df_D_cov <- read_tsv("C:/Users/Dieke/Google Drive/NIOO PhD/AcMNPV_mutationrate/coverage/sub_D_1 (paired, trimmed pairs) mapping.tsv", col_names = FALSE, col_types = cols())
df_E_cov <- read_tsv("C:/Users/Dieke/Google Drive/NIOO PhD/AcMNPV_mutationrate/coverage/sub_E_1 (paired, trimmed pairs) mapping.tsv", col_names = FALSE, col_types = cols())
```

In [4]:

```
clean_and_rank_cov <- function(df){
    filtered_df <- janitor::clean_names(df) %>%
    select(x2, x3, x11, isolate) %>%
    drop_na(x11) %>%
    filter(x3 == "-") %>%
    mutate(rank = percent_rank(desc(x11)))
    return(filtered_df)
}

# clean up dataset; add which isolate is which
df_BAC_cov <- df_BAC_cov %>% mutate(isolate = "BAC") %>% clean_and_rank_cov()
df_A_cov <- df_A_cov %>% mutate(isolate = "A") %>% clean_and_rank_cov()
df_B_cov <- df_B_cov %>% mutate(isolate = "B") %>% clean_and_rank_cov()
df_C_cov <- df_C_cov %>% mutate(isolate = "C") %>% clean_and_rank_cov()
df_D_cov <- df_D_cov %>% mutate(isolate = "D") %>% clean_and_rank_cov()
df_E_cov <- df_E_cov %>% mutate(isolate = "E") %>% clean_and_rank_cov()

# merge datasets into long format table, clean column names using janitor package, select only relevant columns (pos, covg, isolate)
df_cov <- bind_rows(df_BAC_cov, df_A_cov, df_B_cov, df_C_cov, df_D_cov, df_E_cov)
df_cov_clean <- df_cov
```

#### Plotting the coverage across the genome for each isolate¶

In [5]:

```
covg_plot <- ggplot(df_cov_clean, aes(x = x2, y = x11)) +
    geom_line(aes(colour=isolate)) +
    scale_color_brewer(palette = "Dark2") +
    facet_grid(isolate ~ .) +
    labs(title = "Coverage across genome per isolate", x = "Position in genome", y = "Coverage")
#covg_plot
```

Printing the coverage plot, and zooming in so we can see past the weird peak in the "BAC" isolate (from bp 9174-9398). Similar patterns are seen across the different genomes. The peaks in the BAC isolate are a result of remnant plasmid DNA that made up approx. 75% of the reads.

In [6]:

```
covg_plot + theme_light() + coord_cartesian(
  xlim = NULL,
  ylim = c(0, 10000),
  expand = TRUE,
  default = FALSE,
  clip = "on"
)
```

#### Densityplot for coverage per isolate¶

In [7]:

```
density_plot <- ggplot(df_cov_clean, aes(x=x11, fill=isolate)) +
    geom_density(alpha=0.5) +
    #facet_grid(isolate ~ .) +
    scale_fill_brewer(palette = "Dark2") +
    labs(title = "Coverage per isolate", x = "Coverage", y = "Density")

density_plot + theme_light() + coord_cartesian(
  xlim = c(0, 15000),
  expand = TRUE,
  default = FALSE,
  clip = "on"
    )
```

### 3. Filtering mutations¶

#### Read in variant calling data¶

In [8]:

```
df_BAC_raw <- read_csv("C:/Users/Dieke/Google Drive/NIOO PhD/AcMNPV_mutationrate/variants/BAC_H752KDMXX_L1_1.clean (paired, trimmed pairs) mapping variants.csv", col_types = cols())
df_A_raw <- read_csv("C:/Users/Dieke/Google Drive/NIOO PhD/AcMNPV_mutationrate/variants/sub_A_1 (paired, trimmed pairs) mapping variants.csv", col_types = cols())
df_B_raw <- read_csv("C:/Users/Dieke/Google Drive/NIOO PhD/AcMNPV_mutationrate/variants/sub_B_1 (paired, trimmed pairs) mapping variants.csv", col_types = cols())
df_C_raw <- read_csv("C:/Users/Dieke/Google Drive/NIOO PhD/AcMNPV_mutationrate/variants/sub_C_1 (paired, trimmed pairs) mapping variants.csv", col_types = cols())
df_D_raw <- read_csv("C:/Users/Dieke/Google Drive/NIOO PhD/AcMNPV_mutationrate/variants/sub_D_1 (paired, trimmed pairs) mapping variants.csv", col_types = cols())
df_E_raw <- read_csv("C:/Users/Dieke/Google Drive/NIOO PhD/AcMNPV_mutationrate/variants/sub_E_1 (paired, trimmed pairs) mapping variants.csv", col_types = cols())
```

#### Cleaning and merging the variant dataset¶

Adding which isolate is which, and merging datasets into long format table. Column names are cleaned using the Janitor package, mutations are indicated by "nucleotide-position-nucleotide", eg "A295T" for a mutation from A to T at position 295.

In [9]:

```
# cleaning function: add mutations as "nucl-pos-nucl", eg A295T for a mutation from A to T at position 295, clean column names using janitor package
clean_varcall <- function(df){
    filtered_df <- janitor::clean_names(df) %>%
    mutate(mutation = paste(reference,reference_position,allele))
    return(filtered_df)
}

# clean up dataset; add which isolate is which
df_BAC <- df_BAC_raw %>% mutate(isolate = "BAC") %>% clean_varcall()
df_A <- df_A_raw %>% mutate(isolate = "A") %>% clean_varcall()
df_B <- df_B_raw %>% mutate(isolate = "B") %>% clean_varcall()
df_C <- df_C_raw %>% mutate(isolate = "C") %>% clean_varcall()
df_D <- df_D_raw %>% mutate(isolate = "D") %>% clean_varcall() 
df_E <- df_E_raw %>% mutate(isolate = "E") %>% clean_varcall()

# join datasets
df_BAC_full <- left_join(df_BAC, df_BAC_cov, by=c("reference_position" = "x2"))
df_A_full <- left_join(df_A, df_A_cov, by=c("reference_position" = "x2"))
df_B_full <- left_join(df_B, df_B_cov, by=c("reference_position" = "x2"))
df_C_full <- left_join(df_C, df_C_cov, by=c("reference_position" = "x2"))
df_D_full <- left_join(df_D, df_D_cov, by=c("reference_position" = "x2"))
df_E_full <- left_join(df_E, df_E_cov, by=c("reference_position" = "x2"))

# merge datasets into long format table, clean column names using janitor package
df_total <- bind_rows(df_BAC, df_A, df_B, df_C, df_D, df_E)

df_total_full <- bind_rows(df_BAC_full, df_A_full, df_B_full, df_C_full, df_D_full, df_E_full) %>% dplyr::rename(isolate = isolate.y) 
#df_total_full
```

#### Determining which mutations were already there in the ancestral virus population¶

Mutations that have occured multiple times independently are unlikely, therefore I summarise which mutations have a count > 1 and then exclude them.

In [10]:

```
# get mutation counts and add as new column "n"
df_total_clean_counts <- arrange(df_total_full, reference_position) %>% add_count(type, mutation)
```

In [11]:

```
# filtering out mutations which occur more than once
df_singlemutations_unfiltered <- df_total_clean_counts %>% filter(n < 2)
df_singlemutations_unfiltered
```

| reference\_position | type | length | reference | allele | linkage | zygosity | count | coverage | frequency | ... | overlapping\_annotations | coding\_region\_change | amino\_acid\_change | isolate.x | mutation | x3 | x11 | isolate | rank | n |
| --- | --- | --- | --- | --- | --- | --- | --- | --- | --- | --- | --- | --- | --- | --- | --- | --- | --- | --- | --- | --- |
| 11 | SNV | 1 | C | A | NA | Heterozygous | 9 | 1420 | 0.6338028 | ... | NA | NA | NA | E | C 11 A | - | 4330 | E | 0.99289859 | 1 |
| 68 | SNV | 1 | C | T | NA | Heterozygous | 11 | 2154 | 0.5106778 | ... | NA | NA | NA | C | C 68 T | - | 6273 | C | 0.10049222 | 1 |
| 71 | SNV | 1 | T | C | NA | Heterozygous | 13 | 2258 | 0.5757307 | ... | NA | NA | NA | A | T 71 C | - | 6464 | A | 0.09026288 | 1 |
| 132 | SNV | 1 | G | C | NA | Heterozygous | 18 | 2950 | 0.6101695 | ... | NA | NA | NA | E | G 132 C | - | 5480 | E | 0.71205934 | 1 |
| 334 | SNV | 1 | C | A | NA | Heterozygous | 23 | 3696 | 0.6222944 | ... | NA | NA | NA | B | C 334 A | - | 5592 | B | 0.54028488 | 1 |
| 410 | SNV | 1 | A | G | NA | Heterozygous | 21 | 4187 | 0.5015524 | ... | NA | NA | NA | E | A 410 G | - | 5490 | E | 0.70356239 | 1 |
| 1123 | SNV | 1 | A | G | NA | Heterozygous | 242 | 4193 | 5.7715240 | ... | CDS: Ac-bro, Gene: Ac-bro | Ac-bro:c.906T>C | NA | B | A 1123 G | - | 5436 | B | 0.67666227 | 1 |
| 4058 | SNV | 1 | G | T | NA | Heterozygous | 42 | 4635 | 0.9061489 | ... | CDS: Ac-ORF603, Gene: Ac-ORF603 | Ac-ORF603:c.308C>A | Ac-ORF603:p.Pro103His | A | G 4058 T | - | 5608 | A | 0.55243909 | 1 |
| 4473 | SNV | 1 | G | A | NA | Heterozygous | 120 | 4205 | 2.8537455 | ... | Misc. feature: BACMID regioin | NA | NA | E | G 4473 A | - | 5499 | E | 0.69493483 | 1 |
| 5679 | SNV | 1 | G | T | NA | Heterozygous | 15 | 1756 | 0.8542141 | ... | Misc. feature: BACMID regioin | NA | NA | BAC | G 5679 T | - | 2902 | BAC | 0.99604026 | 1 |
| 5680 | SNV | 1 | A | T | NA | Heterozygous | 15 | 2538 | 0.5910165 | ... | Misc. feature: BACMID regioin | NA | NA | E | A 5680 T | - | 3528 | E | 0.99530468 | 1 |
| 5690 | SNV | 1 | A | C | NA | Heterozygous | 13 | 2344 | 0.5546075 | ... | Misc. feature: BACMID regioin | NA | NA | E | A 5690 C | - | 3311 | E | 0.99570340 | 1 |
| 5696 | SNV | 1 | T | G | NA | Heterozygous | 15 | 2322 | 0.6459948 | ... | Misc. feature: BACMID regioin | NA | NA | C | T 5696 G | - | 3233 | C | 0.99575840 | 1 |
| 5700 | Deletion | 1 | T | - | NA | Heterozygous | 16 | 1769 | 0.9044658 | ... | Misc. feature: BACMID regioin | NA | NA | D | T 5700 - | - | 2641 | D | 0.99712644 | 1 |
| 5701 | SNV | 1 | T | G | NA | Heterozygous | 18 | 2028 | 0.8875740 | ... | Misc. feature: BACMID regioin | NA | NA | C | T 5701 G | - | 2841 | C | 0.99640461 | 1 |
| 5703 | Insertion | 1 | - | C | NA | Heterozygous | 39 | 2077 | 1.8777082 | ... | Misc. feature: BACMID regioin | NA | NA | B | - 5703 C | - | 2190 | B | 0.99795826 | 1 |
| 5703 | SNV | 1 | C | G | NA | Homozygous | 10 | 1856 | 0.5387931 | ... | Misc. feature: BACMID regioin | NA | NA | C | C 5703 G | - | 2308 | C | 0.99746329 | 1 |
| 5705 | SNV | 1 | C | A | NA | Heterozygous | 12 | 1723 | 0.6964597 | ... | Misc. feature: BACMID regioin | NA | NA | A | C 5705 A | - | 2193 | A | 0.99792388 | 1 |
| 5707 | SNV | 1 | C | A | NA | Heterozygous | 9 | 1660 | 0.5421687 | ... | Misc. feature: BACMID regioin | NA | NA | B | C 5707 A | - | 2055 | B | 0.99804763 | 1 |
| 5709 | SNV | 1 | C | A | NA | Heterozygous | 18 | 1675 | 1.0746269 | ... | Misc. feature: BACMID regioin | NA | NA | A | C 5709 A | - | 2061 | A | 0.99799263 | 1 |
| 5720 | SNV | 1 | G | T | NA | Heterozygous | 10 | 1415 | 0.7067138 | ... | Misc. feature: BACMID regioin | NA | NA | D | G 5720 T | - | 1684 | D | 0.99847385 | 1 |
| 5726 | SNV | 1 | A | C | NA | Heterozygous | 13 | 1536 | 0.8463542 | ... | Misc. feature: BACMID regioin | NA | NA | C | A 5726 C | - | 1934 | C | 0.99852885 | 1 |
| 5730 | SNV | 1 | T | G | NA | Heterozygous | 16 | 1525 | 1.0491803 | ... | Misc. feature: BACMID regioin | NA | NA | C | T 5730 G | - | 1938 | C | 0.99849447 | 1 |
| 5738 | MNV | 2 | AA | CC | NA | Heterozygous | 16 | 1830 | 0.8743169 | ... | Misc. feature: BACMID regioin | NA | NA | C | AA 5738 CC | - | 1929 | C | 0.99854947 | 1 |
| 5773 | SNV | 1 | T | G | NA | Heterozygous | 17 | 1553 | 1.0946555 | ... | Misc. feature: BACMID regioin | NA | NA | C | T 5773 G | - | 2013 | C | 0.99807513 | 1 |
| 5776 | MNV | 2 | AT | CC | NA | Heterozygous | 9 | 1778 | 0.5061867 | ... | Misc. feature: BACMID regioin | NA | NA | C | AT 5776 CC | - | 2007 | C | 0.99809575 | 1 |
| 5776 | SNV | 1 | A | C | NA | Heterozygous | 12 | 1548 | 0.7751938 | ... | Misc. feature: BACMID regioin | NA | NA | E | A 5776 C | - | 1920 | E | 0.99805450 | 1 |
| 5777 | SNV | 1 | T | A | NA | Heterozygous | 8 | 1416 | 0.5649718 | ... | Misc. feature: BACMID regioin | NA | NA | A | T 5777 A | - | 1840 | A | 0.99828136 | 1 |
| 5804 | SNV | 1 | C | A | NA | Heterozygous | 10 | 1408 | 0.7102273 | ... | Misc. feature: BACMID regioin | NA | NA | E | C 5804 A | - | 1718 | E | 0.99874883 | 1 |
| 5806 | SNV | 1 | A | C | NA | Heterozygous | 7 | 1329 | 0.5267118 | ... | Misc. feature: BACMID regioin | NA | NA | E | A 5806 C | - | 1682 | E | 0.99876945 | 1 |
| ... | ... | ... | ... | ... | ... | ... | ... | ... | ... |  | ... | ... | ... | ... | ... | ... | ... | ... | ... | ... |
| 90082 | SNV | 1 | T | A | NA | Heterozygous | 26 | 4812 | 0.5403159 | ... | CDS: AcOrf-91, Gene: AcOrf-91 | AcOrf-91:c.213A>T | NA | C | T 90082 A | - | 6433 | C | 0.05603448 | 1 |
| 90083 | SNV | 1 | A | C | NA | Heterozygous | 22 | 4320 | 0.5092593 | ... | CDS: AcOrf-91, Gene: AcOrf-91 | AcOrf-91:c.212T>G | AcOrf-91:p.Ile71Arg | C | A 90083 C | - | 6435 | C | 0.05559451 | 1 |
| 90094 | SNV | 1 | A | T | NA | Heterozygous | 33 | 5291 | 0.6237006 | ... | CDS: AcOrf-91, Gene: AcOrf-91 | AcOrf-91:c.201T>A | NA | A | A 90094 T | - | 6971 | A | 0.02440466 | 1 |
| 90106 | SNV | 1 | C | A | NA | Heterozygous | 34 | 4572 | 0.7436570 | ... | CDS: AcOrf-91, Gene: AcOrf-91 | AcOrf-91:c.189G>T | NA | E | C 90106 A | - | 5940 | E | 0.31735687 | 1 |
| 91180 | SNV | 1 | C | T | NA | Heterozygous | 123 | 5590 | 2.2003578 | ... | CDS: AcOrf-93, Gene: AcOrf-93 | AcOrf-93:c.71C>T | AcOrf-93:p.Ala24Val | B | C 91180 T | - | 6269 | B | 0.10538003 | 1 |
| 105200 | MNV | 2 | GA | CT | NA | Heterozygous | 35 | 4925 | 0.7106599 | ... | NA | NA | NA | E | GA 105200 CT | - | 5715 | E | 0.48621652 | 1 |
| 109035 | SNV | 1 | T | C | NA | Heterozygous | 27 | 5287 | 0.5106866 | ... | NA | NA | NA | D | T 109035 C | - | 6075 | D | 0.20340428 | 1 |
| 109042 | SNV | 1 | G | C | NA | Heterozygous | 23 | 4460 | 0.5156951 | ... | NA | NA | NA | B | G 109042 C | - | 5369 | B | 0.73541907 | 1 |
| 109296 | SNV | 1 | G | T | NA | Heterozygous | 33 | 4305 | 0.7665505 | ... | NA | NA | NA | B | G 109296 T | - | 5592 | B | 0.54028488 | 1 |
| 109339 | MNV | 2 | CC | AA | NA | Heterozygous | 25 | 4362 | 0.5731316 | ... | NA | NA | NA | A | CC 109339 AA | - | 5396 | A | 0.74104246 | 1 |
| 109339 | MNV | 3 | CCG | AAT | NA | Heterozygous | 28 | 4417 | 0.6339144 | ... | NA | NA | NA | A | CCG 109339 AAT | - | 5396 | A | 0.74104246 | 1 |
| 109340 | SNV | 1 | C | A | NA | Heterozygous | 23 | 3781 | 0.6083047 | ... | NA | NA | NA | E | C 109340 A | - | 5596 | E | 0.60252021 | 1 |
| 109383 | SNV | 1 | G | A | NA | Heterozygous | 27 | 4291 | 0.6292240 | ... | NA | NA | NA | E | G 109383 A | - | 5822 | E | 0.39567178 | 1 |
| 109385 | SNV | 1 | G | T | NA | Heterozygous | 30 | 4233 | 0.7087172 | ... | NA | NA | NA | E | G 109385 T | - | 5819 | E | 0.39764478 | 1 |
| 109409 | MNV | 2 | TG | CA | NA | Heterozygous | 153 | 4651 | 3.2896151 | ... | NA | NA | NA | A | TG 109409 CA | - | 5397 | A | 0.74010752 | 1 |
| 110855 | SNV | 1 | C | A | NA | Heterozygous | 35 | 4596 | 0.7615318 | ... | CDS: Ac-PIF-3, Gene: Ac-PIF-3 | Ac-PIF-3:c.515G>T | Ac-PIF-3:p.Gly172Val | B | C 110855 A | - | 5098 | B | 0.89017901 | 1 |
| 117608 | SNV | 1 | G | A | NA | Heterozygous | 88 | 4994 | 1.7621145 | ... | CDS: Ac-ChiA, Gene: Ac-ChiA | Ac-ChiA:c.903C>T | NA | D | G 117608 A | - | 5604 | D | 0.58751306 | 1 |
| 126306 | SNV | 1 | T | C | NA | Heterozygous | 271 | 5133 | 5.2795636 | ... | CDS: Ac-p94, Gene: Ac-p94 | Ac-p94:c.1550A>G | Ac-p94:p.Gln517Arg | D | T 126306 C | - | 5895 | D | 0.34204339 | 1 |
| 129170 | SNV | 1 | C | A | NA | Heterozygous | 29 | 4672 | 0.6207192 | ... | NA | NA | NA | E | C 129170 A | - | 6017 | E | 0.26794258 | 1 |
| 129180 | SNV | 1 | G | A | NA | Heterozygous | 33 | 4583 | 0.7200524 | ... | NA | NA | NA | E | G 129180 A | - | 5979 | E | 0.29242974 | 1 |
| 129266 | SNV | 1 | G | T | NA | Heterozygous | 29 | 4573 | 0.6341570 | ... | NA | NA | NA | B | G 129266 T | - | 5812 | B | 0.34662872 | 1 |
| 129272 | SNV | 1 | C | A | NA | Heterozygous | 36 | 4453 | 0.8084437 | ... | NA | NA | NA | B | C 129272 A | - | 5798 | B | 0.35848045 | 1 |
| 129273 | SNV | 1 | G | T | NA | Heterozygous | 52 | 4609 | 1.1282274 | ... | NA | NA | NA | A | G 129273 T | - | 6383 | A | 0.11182836 | 1 |
| 129359 | SNV | 1 | G | T | NA | Heterozygous | 41 | 4477 | 0.9157918 | ... | NA | NA | NA | C | G 129359 T | - | 5862 | C | 0.35950476 | 1 |
| 129464 | MNV | 2 | AA | GG | NA | Heterozygous | 30 | 5043 | 0.5948840 | ... | NA | NA | NA | D | AA 129464 GG | - | 5937 | D | 0.30678381 | 1 |
| 129554 | SNV | 1 | C | T | NA | Heterozygous | 22 | 4374 | 0.5029721 | ... | NA | NA | NA | A | C 129554 T | - | 5470 | A | 0.67600918 | 1 |
| 144395 | SNV | 1 | C | T | NA | Heterozygous | 373 | 4751 | 7.8509787 | ... | CDS: Ac-pe38, Gene: Ac-pe38 | Ac-pe38:c.299C>T | Ac-pe38:p.Thr100Ile | E | C 144395 T | - | 5345 | E | 0.81810620 | 1 |
| 145127 | Insertion | 1 | - | T | NA | Heterozygous | 28 | 5320 | 0.5263158 | ... | NA | NA | NA | D | - 145127 T | - | 5971 | D | 0.27903124 | 1 |
| 145440 | SNV | 1 | T | C | NA | Heterozygous | 13 | 1877 | 0.6925946 | ... | NA | NA | NA | E | T 145440 C | - | 3786 | E | 0.99472722 | 1 |
| 145447 | SNV | 1 | C | T | NA | Heterozygous | 14 | 1695 | 0.8259587 | ... | NA | NA | NA | E | C 145447 T | - | 3540 | E | 0.99527718 | 1 |

#### Filter variants¶

The function **rules\_filtering** lists the criterea for inclusion of a variant in the downstream variant calling, which is as follows:

- forward reverse balance > 0.05
- count > 10
- type is "SNV"
- frequency is > 0.5 | 1 | 2 %
- coverage is within the 98% of ranked covg (i.e. not in the extreme ends of the tails of the distribution.

In [12]:

```
rules_filtering <- function(df){
  filtered_df <- filter(df,
         forward_reverse_balance > 0.05,
         count > 10,
         type == "SNV",
         frequency > 0.5,
         rank >= 0.01,
         rank <= 0.99
                       )
  return(filtered_df)
}
```

In [13]:

```
# filtering variants for whole genome
df_singlemutations <- df_singlemutations_unfiltered %>%
  rules_filtering() 

df_singlemutations
```

| reference\_position | type | length | reference | allele | linkage | zygosity | count | coverage | frequency | ... | overlapping\_annotations | coding\_region\_change | amino\_acid\_change | isolate.x | mutation | x3 | x11 | isolate | rank | n |
| --- | --- | --- | --- | --- | --- | --- | --- | --- | --- | --- | --- | --- | --- | --- | --- | --- | --- | --- | --- | --- |
| 68 | SNV | 1 | C | T | NA | Heterozygous | 11 | 2154 | 0.5106778 | ... | NA | NA | NA | C | C 68 T | - | 6273 | C | 0.10049222 | 1 |
| 71 | SNV | 1 | T | C | NA | Heterozygous | 13 | 2258 | 0.5757307 | ... | NA | NA | NA | A | T 71 C | - | 6464 | A | 0.09026288 | 1 |
| 132 | SNV | 1 | G | C | NA | Heterozygous | 18 | 2950 | 0.6101695 | ... | NA | NA | NA | E | G 132 C | - | 5480 | E | 0.71205934 | 1 |
| 334 | SNV | 1 | C | A | NA | Heterozygous | 23 | 3696 | 0.6222944 | ... | NA | NA | NA | B | C 334 A | - | 5592 | B | 0.54028488 | 1 |
| 410 | SNV | 1 | A | G | NA | Heterozygous | 21 | 4187 | 0.5015524 | ... | NA | NA | NA | E | A 410 G | - | 5490 | E | 0.70356239 | 1 |
| 1123 | SNV | 1 | A | G | NA | Heterozygous | 242 | 4193 | 5.7715240 | ... | CDS: Ac-bro, Gene: Ac-bro | Ac-bro:c.906T>C | NA | B | A 1123 G | - | 5436 | B | 0.67666227 | 1 |
| 4058 | SNV | 1 | G | T | NA | Heterozygous | 42 | 4635 | 0.9061489 | ... | CDS: Ac-ORF603, Gene: Ac-ORF603 | Ac-ORF603:c.308C>A | Ac-ORF603:p.Pro103His | A | G 4058 T | - | 5608 | A | 0.55243909 | 1 |
| 4473 | SNV | 1 | G | A | NA | Heterozygous | 120 | 4205 | 2.8537455 | ... | Misc. feature: BACMID regioin | NA | NA | E | G 4473 A | - | 5499 | E | 0.69493483 | 1 |
| 7383 | SNV | 1 | C | A | NA | Heterozygous | 37 | 3548 | 1.0428410 | ... | Misc. feature: BACMID regioin | NA | NA | D | C 7383 A | - | 4584 | D | 0.98400979 | 1 |
| 7400 | SNV | 1 | G | T | NA | Heterozygous | 45 | 4162 | 1.0812110 | ... | Misc. feature: BACMID regioin | NA | NA | C | G 7400 T | - | 5210 | C | 0.91181323 | 1 |
| 9990 | SNV | 1 | A | G | NA | Heterozygous | 18 | 3098 | 0.5810200 | ... | Misc. feature: BACMID regioin | NA | NA | B | A 9990 G | - | 3895 | B | 0.98882885 | 1 |
| 10035 | SNV | 1 | C | T | NA | Heterozygous | 23 | 2958 | 0.7775524 | ... | Misc. feature: BACMID regioin | NA | NA | BAC | C 10035 T | - | 4984 | BAC | 0.89459935 | 1 |
| 10065 | SNV | 1 | A | T | NA | Heterozygous | 16 | 2583 | 0.6194348 | ... | Misc. feature: BACMID regioin | NA | NA | E | A 10065 T | - | 4634 | E | 0.98803140 | 1 |
| 10353 | SNV | 1 | T | G | NA | Heterozygous | 18 | 2992 | 0.6016043 | ... | Misc. feature: BACMID regioin | NA | NA | A | T 10353 G | - | 4861 | A | 0.96655530 | 1 |
| 10392 | SNV | 1 | G | A | NA | Heterozygous | 15 | 2755 | 0.5444646 | ... | Misc. feature: BACMID regioin | NA | NA | B | G 10392 A | - | 4106 | B | 0.97932134 | 1 |
| 11404 | SNV | 1 | A | C | NA | Heterozygous | 29 | 4202 | 0.6901475 | ... | Misc. feature: BACMID regioin | NA | NA | A | A 11404 C | - | 5771 | A | 0.42537673 | 1 |
| 23894 | SNV | 1 | C | T | NA | Heterozygous | 81 | 4747 | 1.7063408 | ... | CDS: Ac-egt, Gene: Ac-egt | Ac-egt:c.836C>T | Ac-egt:p.Pro279Leu | D | C 23894 T | - | 5450 | D | 0.70882830 | 1 |
| 37958 | SNV | 1 | T | C | NA | Heterozygous | 26 | 4545 | 0.5720572 | ... | NA | NA | NA | A | T 37958 C | - | 5606 | A | 0.55437772 | 1 |
| 38215 | SNV | 1 | G | A | NA | Heterozygous | 24 | 3458 | 0.6940428 | ... | NA | NA | NA | A | G 38215 A | - | 5558 | A | 0.59651185 | 1 |
| 38235 | SNV | 1 | G | T | NA | Heterozygous | 43 | 3230 | 1.3312693 | ... | NA | NA | NA | A | G 38235 T | - | 5760 | A | 0.43324121 | 1 |
| 38241 | SNV | 1 | G | T | NA | Heterozygous | 44 | 3251 | 1.3534297 | ... | NA | NA | NA | A | G 38241 T | - | 5778 | A | 0.42029643 | 1 |
| 38242 | SNV | 1 | G | T | NA | Heterozygous | 27 | 3307 | 0.8164500 | ... | NA | NA | NA | E | G 38242 T | - | 5724 | E | 0.47758896 | 1 |
| 38245 | SNV | 1 | C | A | NA | Heterozygous | 41 | 3299 | 1.2428008 | ... | NA | NA | NA | A | C 38245 A | - | 5867 | A | 0.35702992 | 1 |
| 38256 | SNV | 1 | C | G | NA | Heterozygous | 22 | 3532 | 0.6228766 | ... | NA | NA | NA | A | C 38256 G | - | 5985 | A | 0.27757383 | 1 |
| 38349 | SNV | 1 | A | G | NA | Heterozygous | 28 | 3633 | 0.7707129 | ... | NA | NA | NA | E | A 38349 G | - | 6683 | E | 0.03022741 | 1 |
| 38399 | SNV | 1 | T | A | NA | Heterozygous | 15 | 2293 | 0.6541648 | ... | NA | NA | NA | C | T 38399 A | - | 5203 | C | 0.91452181 | 1 |
| 38400 | SNV | 1 | G | T | NA | Heterozygous | 27 | 2184 | 1.2362637 | ... | NA | NA | NA | A | G 38400 T | - | 5144 | A | 0.89196640 | 1 |
| 38440 | SNV | 1 | A | T | NA | Heterozygous | 14 | 2783 | 0.5030543 | ... | NA | NA | NA | C | A 38440 T | - | 5393 | C | 0.80055546 | 1 |
| 40156 | SNV | 1 | G | A | NA | Heterozygous | 92 | 4844 | 1.8992568 | ... | CDS: AcOrf-34, Gene: AcOrf-34 | AcOrf-34:c.420C>T | NA | D | G 40156 A | - | 5464 | D | 0.69845460 | 1 |
| 49063 | SNV | 1 | A | C | NA | Heterozygous | 33 | 4165 | 0.7923169 | ... | CDS: Ac-odv-e66, Gene: Ac-odv-e66 | Ac-odv-e66:c.712A>C | Ac-odv-e66:p.Met238Leu | A | A 49063 C | - | 5827 | A | 0.38542182 | 1 |
| ... | ... | ... | ... | ... | ... | ... | ... | ... | ... |  | ... | ... | ... | ... | ... | ... | ... | ... | ... | ... |
| 82741 | SNV | 1 | C | T | NA | Heterozygous | 21 | 3963 | 0.5299016 | ... | NA | NA | NA | C | C 82741 T | - | 5591 | C | 0.62076528 | 1 |
| 82775 | SNV | 1 | G | T | NA | Heterozygous | 27 | 4500 | 0.6000000 | ... | NA | NA | NA | E | G 82775 T | - | 5575 | E | 0.62213331 | 1 |
| 82777 | SNV | 1 | A | G | NA | Heterozygous | 26 | 4557 | 0.5705508 | ... | NA | NA | NA | E | A 82777 G | - | 5575 | E | 0.62213331 | 1 |
| 89888 | SNV | 1 | C | A | NA | Heterozygous | 43 | 5182 | 0.8297954 | ... | CDS: AcOrf-91, Gene: AcOrf-91 | AcOrf-91:c.407G>T | AcOrf-91:p.Ser136Ile | B | C 89888 A | - | 5941 | B | 0.25153303 | 1 |
| 89941 | SNV | 1 | C | A | NA | Heterozygous | 68 | 4658 | 1.4598540 | ... | CDS: AcOrf-91, Gene: AcOrf-91 | AcOrf-91:c.354G>T | NA | B | C 89941 A | - | 5810 | B | 0.34812737 | 1 |
| 90012 | SNV | 1 | A | T | NA | Heterozygous | 43 | 4725 | 0.9100529 | ... | CDS: AcOrf-91, Gene: AcOrf-91 | AcOrf-91:c.283T>A | AcOrf-91:p.Ser95Thr | E | A 90012 T | - | 6334 | E | 0.10157152 | 1 |
| 90063 | SNV | 1 | T | A | NA | Heterozygous | 27 | 4677 | 0.5772931 | ... | CDS: AcOrf-91, Gene: AcOrf-91 | AcOrf-91:c.232A>T | AcOrf-91:p.Thr78Ser | E | T 90063 A | - | 6075 | E | 0.23218803 | 1 |
| 90078 | SNV | 1 | G | A | NA | Heterozygous | 41 | 5516 | 0.7432922 | ... | CDS: AcOrf-91, Gene: AcOrf-91 | AcOrf-91:c.217C>T | AcOrf-91:p.Pro73Ser | A | G 90078 A | - | 6967 | A | 0.02455590 | 1 |
| 90082 | SNV | 1 | T | A | NA | Heterozygous | 26 | 4812 | 0.5403159 | ... | CDS: AcOrf-91, Gene: AcOrf-91 | AcOrf-91:c.213A>T | NA | C | T 90082 A | - | 6433 | C | 0.05603448 | 1 |
| 90083 | SNV | 1 | A | C | NA | Heterozygous | 22 | 4320 | 0.5092593 | ... | CDS: AcOrf-91, Gene: AcOrf-91 | AcOrf-91:c.212T>G | AcOrf-91:p.Ile71Arg | C | A 90083 C | - | 6435 | C | 0.05559451 | 1 |
| 90094 | SNV | 1 | A | T | NA | Heterozygous | 33 | 5291 | 0.6237006 | ... | CDS: AcOrf-91, Gene: AcOrf-91 | AcOrf-91:c.201T>A | NA | A | A 90094 T | - | 6971 | A | 0.02440466 | 1 |
| 90106 | SNV | 1 | C | A | NA | Heterozygous | 34 | 4572 | 0.7436570 | ... | CDS: AcOrf-91, Gene: AcOrf-91 | AcOrf-91:c.189G>T | NA | E | C 90106 A | - | 5940 | E | 0.31735687 | 1 |
| 91180 | SNV | 1 | C | T | NA | Heterozygous | 123 | 5590 | 2.2003578 | ... | CDS: AcOrf-93, Gene: AcOrf-93 | AcOrf-93:c.71C>T | AcOrf-93:p.Ala24Val | B | C 91180 T | - | 6269 | B | 0.10538003 | 1 |
| 109035 | SNV | 1 | T | C | NA | Heterozygous | 27 | 5287 | 0.5106866 | ... | NA | NA | NA | D | T 109035 C | - | 6075 | D | 0.20340428 | 1 |
| 109042 | SNV | 1 | G | C | NA | Heterozygous | 23 | 4460 | 0.5156951 | ... | NA | NA | NA | B | G 109042 C | - | 5369 | B | 0.73541907 | 1 |
| 109296 | SNV | 1 | G | T | NA | Heterozygous | 33 | 4305 | 0.7665505 | ... | NA | NA | NA | B | G 109296 T | - | 5592 | B | 0.54028488 | 1 |
| 109340 | SNV | 1 | C | A | NA | Heterozygous | 23 | 3781 | 0.6083047 | ... | NA | NA | NA | E | C 109340 A | - | 5596 | E | 0.60252021 | 1 |
| 109383 | SNV | 1 | G | A | NA | Heterozygous | 27 | 4291 | 0.6292240 | ... | NA | NA | NA | E | G 109383 A | - | 5822 | E | 0.39567178 | 1 |
| 109385 | SNV | 1 | G | T | NA | Heterozygous | 30 | 4233 | 0.7087172 | ... | NA | NA | NA | E | G 109385 T | - | 5819 | E | 0.39764478 | 1 |
| 110855 | SNV | 1 | C | A | NA | Heterozygous | 35 | 4596 | 0.7615318 | ... | CDS: Ac-PIF-3, Gene: Ac-PIF-3 | Ac-PIF-3:c.515G>T | Ac-PIF-3:p.Gly172Val | B | C 110855 A | - | 5098 | B | 0.89017901 | 1 |
| 117608 | SNV | 1 | G | A | NA | Heterozygous | 88 | 4994 | 1.7621145 | ... | CDS: Ac-ChiA, Gene: Ac-ChiA | Ac-ChiA:c.903C>T | NA | D | G 117608 A | - | 5604 | D | 0.58751306 | 1 |
| 126306 | SNV | 1 | T | C | NA | Heterozygous | 271 | 5133 | 5.2795636 | ... | CDS: Ac-p94, Gene: Ac-p94 | Ac-p94:c.1550A>G | Ac-p94:p.Gln517Arg | D | T 126306 C | - | 5895 | D | 0.34204339 | 1 |
| 129170 | SNV | 1 | C | A | NA | Heterozygous | 29 | 4672 | 0.6207192 | ... | NA | NA | NA | E | C 129170 A | - | 6017 | E | 0.26794258 | 1 |
| 129180 | SNV | 1 | G | A | NA | Heterozygous | 33 | 4583 | 0.7200524 | ... | NA | NA | NA | E | G 129180 A | - | 5979 | E | 0.29242974 | 1 |
| 129266 | SNV | 1 | G | T | NA | Heterozygous | 29 | 4573 | 0.6341570 | ... | NA | NA | NA | B | G 129266 T | - | 5812 | B | 0.34662872 | 1 |
| 129272 | SNV | 1 | C | A | NA | Heterozygous | 36 | 4453 | 0.8084437 | ... | NA | NA | NA | B | C 129272 A | - | 5798 | B | 0.35848045 | 1 |
| 129273 | SNV | 1 | G | T | NA | Heterozygous | 52 | 4609 | 1.1282274 | ... | NA | NA | NA | A | G 129273 T | - | 6383 | A | 0.11182836 | 1 |
| 129359 | SNV | 1 | G | T | NA | Heterozygous | 41 | 4477 | 0.9157918 | ... | NA | NA | NA | C | G 129359 T | - | 5862 | C | 0.35950476 | 1 |
| 129554 | SNV | 1 | C | T | NA | Heterozygous | 22 | 4374 | 0.5029721 | ... | NA | NA | NA | A | C 129554 T | - | 5470 | A | 0.67600918 | 1 |
| 144395 | SNV | 1 | C | T | NA | Heterozygous | 373 | 4751 | 7.8509787 | ... | CDS: Ac-pe38, Gene: Ac-pe38 | Ac-pe38:c.299C>T | Ac-pe38:p.Thr100Ile | E | C 144395 T | - | 5345 | E | 0.81810620 | 1 |

#### The number of mutations that are "left" in the whole genome after the filtering steps are summarised below¶

In [14]:

```
df_summary <- df_singlemutations %>% group_by(isolate) %>% dplyr::summarize(count = n())
df_summary
```

| isolate | count |
| --- | --- |
| A | 20 |
| B | 14 |
| BAC | 1 |
| C | 9 |
| D | 7 |
| E | 18 |

#### We can do the same for only the BACmid region, which results in the following mutation counts¶

In [15]:

```
# filtering variants for bacmid region only
df_singlemutations_BACMID <- df_singlemutations %>%
  filter(overlapping_annotations == "Misc. feature: BACMID regioin") #filters for only bacmid region
#df_singlemutations_BACMID

df_summary_BACMID <- df_singlemutations_BACMID %>% group_by(isolate) %>% dplyr::summarize(count = n())
df_summary_BACMID
```

| isolate | count |
| --- | --- |
| A | 2 |
| B | 2 |
| BAC | 1 |
| C | 1 |
| D | 1 |
| E | 2 |

### 4. Plot mutations in the coverage plot¶

Merging the datasets (coverage and variant calling) so the mutations can be plotted along the coverageplots.

In [16]:

```
merged_plot_data <- left_join(df_cov_clean, df_singlemutations, by = c("x2" = "reference_position", "isolate")) %>% 
    dplyr::rename(x11 = x11.x) %>%
    mutate(covg = case_when(type == "SNV" ~ x11))
```

In [17]:

```
covg_variants_plot <- ggplot(merged_plot_data, aes(x = x2, y = x11)) +
    geom_line(aes(colour=isolate)) +
    scale_color_brewer(palette = "Dark2") +
    facet_grid(isolate ~ .) +
    geom_point(aes(x = x2, y = covg)) +
    labs(title = "Coverage across genome per isolate; mutations indicated by a dot", x = "Position in genome", y = "Coverage")

covg_variants_plot + theme_light() + coord_cartesian(
  xlim = c(0, 150000),
  ylim = c(0, 10000),
  expand = TRUE,
  default = FALSE,
  clip = "on"
    )
```

```
Warning message:
"Removed 872721 rows containing missing values (geom_point)."
```

In [ ]:
