## Supplementary material for "Empirical estimates of the mutation rate for an alphabaculovirus": Appendix 3 Subsampling MutationCalling v3 freq=1.html

In [12]:

```
rules_filtering <- function(df){
  filtered_df <- filter(df,
         forward_reverse_balance > 0.05,
         count > 10,
         type == "SNV",
         frequency > 1,
         rank >= 0.01,
         rank <= 0.99
                       )
  return(filtered_df)
}
```

In [13]:

```
# filtering variants for whole genome
df_singlemutations <- df_singlemutations_unfiltered %>%
  rules_filtering() 

df_singlemutations
```

| reference\_position | type | length | reference | allele | linkage | zygosity | count | coverage | frequency | ... | overlapping\_annotations | coding\_region\_change | amino\_acid\_change | isolate.x | mutation | x3 | x11 | isolate | rank | n |
| --- | --- | --- | --- | --- | --- | --- | --- | --- | --- | --- | --- | --- | --- | --- | --- | --- | --- | --- | --- | --- |
| 1123 | SNV | 1 | A | G | NA | Heterozygous | 242 | 4193 | 5.771524 | ... | CDS: Ac-bro, Gene: Ac-bro | Ac-bro:c.906T>C | NA | B | A 1123 G | - | 5436 | B | 0.6766623 | 1 |
| 4473 | SNV | 1 | G | A | NA | Heterozygous | 120 | 4205 | 2.853746 | ... | Misc. feature: BACMID regioin | NA | NA | E | G 4473 A | - | 5499 | E | 0.6949348 | 1 |
| 7383 | SNV | 1 | C | A | NA | Heterozygous | 37 | 3548 | 1.042841 | ... | Misc. feature: BACMID regioin | NA | NA | D | C 7383 A | - | 4584 | D | 0.9840098 | 1 |
| 7400 | SNV | 1 | G | T | NA | Heterozygous | 45 | 4162 | 1.081211 | ... | Misc. feature: BACMID regioin | NA | NA | C | G 7400 T | - | 5210 | C | 0.9118132 | 1 |
| 23894 | SNV | 1 | C | T | NA | Heterozygous | 81 | 4747 | 1.706341 | ... | CDS: Ac-egt, Gene: Ac-egt | Ac-egt:c.836C>T | Ac-egt:p.Pro279Leu | D | C 23894 T | - | 5450 | D | 0.7088283 | 1 |
| 38235 | SNV | 1 | G | T | NA | Heterozygous | 43 | 3230 | 1.331269 | ... | NA | NA | NA | A | G 38235 T | - | 5760 | A | 0.4332412 | 1 |
| 38241 | SNV | 1 | G | T | NA | Heterozygous | 44 | 3251 | 1.353430 | ... | NA | NA | NA | A | G 38241 T | - | 5778 | A | 0.4202964 | 1 |
| 38245 | SNV | 1 | C | A | NA | Heterozygous | 41 | 3299 | 1.242801 | ... | NA | NA | NA | A | C 38245 A | - | 5867 | A | 0.3570299 | 1 |
| 38400 | SNV | 1 | G | T | NA | Heterozygous | 27 | 2184 | 1.236264 | ... | NA | NA | NA | A | G 38400 T | - | 5144 | A | 0.8919664 | 1 |
| 40156 | SNV | 1 | G | A | NA | Heterozygous | 92 | 4844 | 1.899257 | ... | CDS: AcOrf-34, Gene: AcOrf-34 | AcOrf-34:c.420C>T | NA | D | G 40156 A | - | 5464 | D | 0.6984546 | 1 |
| 50202 | SNV | 1 | C | A | NA | Heterozygous | 124 | 5321 | 2.330389 | ... | CDS: Ac-odv-e66, Gene: Ac-odv-e66 | Ac-odv-e66:c.1851C>A | NA | A | C 50202 A | - | 6183 | A | 0.1806564 | 1 |
| 52396 | SNV | 1 | G | T | NA | Heterozygous | 979 | 4329 | 22.614923 | ... | CDS: Ac-lef8, Gene: Ac-lef8 | Ac-lef8:c.2392C>A | Ac-lef8:p.Leu798Ile | D | G 52396 T | - | 4926 | D | 0.9601963 | 1 |
| 75791 | SNV | 1 | G | A | NA | Heterozygous | 554 | 5273 | 10.506353 | ... | CDS: Ac-vlf-1, Gene: Ac-vlf-1 | Ac-vlf-1:c.798C>T | NA | C | G 75791 A | - | 5949 | C | 0.2839534 | 1 |
| 82741 | SNV | 1 | C | A | NA | Heterozygous | 45 | 3833 | 1.174015 | ... | NA | NA | NA | A | C 82741 A | - | 5713 | A | 0.4688926 | 1 |
| 89941 | SNV | 1 | C | A | NA | Heterozygous | 68 | 4658 | 1.459854 | ... | CDS: AcOrf-91, Gene: AcOrf-91 | AcOrf-91:c.354G>T | NA | B | C 89941 A | - | 5810 | B | 0.3481274 | 1 |
| 91180 | SNV | 1 | C | T | NA | Heterozygous | 123 | 5590 | 2.200358 | ... | CDS: AcOrf-93, Gene: AcOrf-93 | AcOrf-93:c.71C>T | AcOrf-93:p.Ala24Val | B | C 91180 T | - | 6269 | B | 0.1053800 | 1 |
| 117608 | SNV | 1 | G | A | NA | Heterozygous | 88 | 4994 | 1.762115 | ... | CDS: Ac-ChiA, Gene: Ac-ChiA | Ac-ChiA:c.903C>T | NA | D | G 117608 A | - | 5604 | D | 0.5875131 | 1 |
| 126306 | SNV | 1 | T | C | NA | Heterozygous | 271 | 5133 | 5.279564 | ... | CDS: Ac-p94, Gene: Ac-p94 | Ac-p94:c.1550A>G | Ac-p94:p.Gln517Arg | D | T 126306 C | - | 5895 | D | 0.3420434 | 1 |
| 129273 | SNV | 1 | G | T | NA | Heterozygous | 52 | 4609 | 1.128227 | ... | NA | NA | NA | A | G 129273 T | - | 6383 | A | 0.1118284 | 1 |
| 144395 | SNV | 1 | C | T | NA | Heterozygous | 373 | 4751 | 7.850979 | ... | CDS: Ac-pe38, Gene: Ac-pe38 | Ac-pe38:c.299C>T | Ac-pe38:p.Thr100Ile | E | C 144395 T | - | 5345 | E | 0.8181062 | 1 |

#### The number of mutations that are "left" in the whole genome after the filtering steps are summarised below¶

In [14]:

```
df_summary <- df_singlemutations %>% group_by(isolate) %>% dplyr::summarize(count = n())
df_summary
```

| isolate | count |
| --- | --- |
| A | 7 |
| B | 3 |
| C | 2 |
| D | 6 |
| E | 2 |

#### We can do the same for only the BACmid region, which results in the following mutation counts¶

In [15]:

```
# filtering variants for bacmid region only
df_singlemutations_BACMID <- df_singlemutations %>%
  filter(overlapping_annotations == "Misc. feature: BACMID regioin") #filters for only bacmid region
#df_singlemutations_BACMID

df_summary_BACMID <- df_singlemutations_BACMID %>% group_by(isolate) %>% dplyr::summarize(count = n())
df_summary_BACMID
```

| isolate | count |
| --- | --- |
| C | 1 |
| D | 1 |
| E | 1 |

### 4. Plot mutations in the coverage plot¶

Merging the datasets (coverage and variant calling) so the mutations can be plotted along the coverageplots.

In [16]:

```
merged_plot_data <- left_join(df_cov_clean, df_singlemutations, by = c("x2" = "reference_position", "isolate")) %>% 
    dplyr::rename(x11 = x11.x) %>%
    mutate(covg = case_when(type == "SNV" ~ x11))
```

In [17]:

```
covg_variants_plot <- ggplot(merged_plot_data, aes(x = x2, y = x11)) +
    geom_line(aes(colour=isolate)) +
    scale_color_brewer(palette = "Dark2") +
    facet_grid(isolate ~ .) +
    geom_point(aes(x = x2, y = covg)) +
    labs(title = "Coverage across genome per isolate; mutations indicated by a dot", x = "Position in genome", y = "Coverage")

covg_variants_plot + theme_light() + coord_cartesian(
  xlim = c(0, 150000),
  ylim = c(0, 10000),
  expand = TRUE,
  default = FALSE,
  clip = "on"
    )
```

```
Warning message:
"Removed 872770 rows containing missing values (geom_point)."
```

In [ ]:
