## Supplementary material for "Empirical estimates of the mutation rate for an alphabaculovirus": Appendix 5 Model mutation rates bacmid.txt

#########################################################################################################
#                                                                                                       #
# Authors: Mark Zwart & Dieke Boezen                                                                   	#
# Date: April 19, 2018; February 21, 2020; March 3, 2020, September 30, 2020, May 10, 2021		#
#                                                                                                    	#
# The final version of a model used to predict the mutation rate based on the frequency of	        #
# mutations in neutral locus in an evolved popuation. This is the final version used to estimte         #
# the mutation rate for the bacmid data.								#
#                                                                                                    	#
#########################################################################################################    

# Note: the model does not include linkage between loci.

# Defining single function for bottlenecking and passaging the virus population once in a single
# animal

bottleneck.evolve <- function(vir.pop, bottleneck.size, final.size, virus.replication, mutation.rate)

{
	# First determine a few parameters. Some are parameters that are defined for this model, but 
	# are included here to make the function more general.
	
	g.size = length(vir.pop)
	
	# Infection bottleneck: draw a sample from the existing population. Draw the number of mutants 
	# from a binomial distribution. First draw the number of founders from a Poisson distribution, 
	# which is also the initial pop size. Then draw mutants that pass through bottleneck for each 
	# genome position, and the rest of the population is per definition wildtype at each position.
	vir.pop.size = rpois(n = 1, lambda = bottleneck.size) 
	mt = rbinom(n = g.size, size = vir.pop.size, prob = vir.pop)
	wt = vir.pop.size - mt
		
	# Let the population grow and mutate until the final size is reached.
	
	while(vir.pop.size < final.size) {
		
		# In the final round of replication adjust progeny size to exactly reach final.size
		if(vir.pop.size + (vir.pop.size*virus.replication) < final.size) {
			progeny.now = vir.pop.size*virus.replication
		} else {
			progeny.now = final.size - vir.pop.size
		} # end of if else statement.
		
		# Determine new population size (just for counter)
		vir.pop.size = vir.pop.size + progeny.now
		
		# Divide progeny over mt and wt
		freq.mt = mt/(mt+wt)                # Frequency of the mutants
		new.mt = round(freq.mt*progeny.now) # Round because we are modeling absolute numbers
		new.wt = progeny.now - new.mt       # That which is not mutated is wt, for now
		
		if(progeny.now*max.stoch > 1/mutation.rate) denovo.mt = new.wt*mutation.rate else {
			denovo.mt = rbinom(n = g.size, size = new.wt, prob = mutation.rate)}
		mt = mt + new.mt + denovo.mt
		wt = wt + new.wt - denovo.mt
		
	} # end of while loop for pop growth. 
	
	vir.pop = mt/(mt+wt)
	return(vir.pop)	
}


#########################################################################################################

# Notes on model parameter estimates:

# Bottleneck size estimates: we start with the relationship between survival (S) after exporsure
# to single dose vs. the founding population size (Lambda), which following Zwart et al. 2009 Proc. R. 
# Soc. B (2009) 276, 2233�2242 can be approximated as 1-M = S = exp(-Lambda). So for an 10 X LD_99 dose,   
# the corresponding 10*-ln(1-0.99) ~ 46.

# Final size: We counted OBs after each round of passaging for the pool of five insects, leading to an 
# mean yield of 1.33*10^6 OBs/larva. We are not aware of a good estimate of the number of occlusion 
# derived viruses per occlusion body, so we take a value of 100 as an approximation. From Zwart et al. 
# 2008 J Virol Methods 148:146-154 we know that there are ~4 nucleocapsids per OB. Hence the yield (in
# genomes) per infected larva becomes 1.33*10^6 * 100 * 4 = 5.32*10^8 genomes/larva.

# Genome size: for the whole genome, the reference used by Dieke for the bioinformatics has a length
# of the genome (with the bacmid inserted) is 145,465 bp. The complete length of the bacmid reference
# is 12,510 bp, but we need to subtract the length of (1) the intact polyhedrin in FastBac-Dual (-739 bp)
# and its promotor sequence (-126 bp). The other bits (disrupted polyhedrin fragements from the ancestral
# copy and the P10 promotor in pFastBac-Dual are considered neutral. So this renders a neutral insert
# with a size of 11,646 bp.

#########################################################################################################


# Basic model parameters that are not explored. Names as given the manuscript included in each description
# in parentheses.

bottleneck.size <-	46		# The number of virus founders i.e., genomes effectively 
					# initiating infection per insect. (Lambda)
final.size <- 		5.32*10^8	# The number of virus individuals coming from a dead larvae. (Kappa)
number.passages <- 	5		# The number of serial passages of infection the virus 
					# population has undergone.
genome.size <- 		11646         	# Size of our model genome. Note that we only consider the 
					# neutral bacmid insert, in terms of genome size in the 
					# simulation and mutations in the experimental data. Not that 
					# for a low mutation rate and large genome size this model gives
					# a reasonable approx. It will be useless at predictions for 
					# small genome sizes and high mutation rates. (g)
max.stoch <-		10000		# When population size * max.stoch > mutation rate^-1, mutation 
					# switches from being stochastic to being deterministic. (Phi)
replicates <-		1000		# Number of independent simulations to determine the number of 
					# mutations per experimental replicate. Note that number of
					# replicates has to be large in order for this code to function 
					# properly. Our recommendation is 1000 replicates, an absolute
					# minimum would be ~ 100. 
max.sim.data <-		1000		# Parameter value that only determines the size and max value
					# mutations per genome. This value can be exceeded in the
					# simulations, but these results are then removed in subsequent
					# model fitting. Since the number of categories with a prediction 
					# of frequency zero and zero observations does not affect the
					# multinomial likelihood, it is not a problem to set this to a 
					# high value. However, note that if the code is being used to 
					# predict data with a large number of mutations per replicate
					# (i.e., >> 10), you really need a very large number of replicate
					# simulations to get a good pseudolikelihood approximation.
					# (max.sim.data + 1 = sigma)
detect.thresholds <- c(0.01, 0.005, 0.02)
					# The frequency mutations must reach in order to be included in
					# the analysis. 0.01 is probably the best value for this
					# analysis, but some additional values with a two-fold difference
					# are included in this version so that one does not have to 
					# repeat the simulations to analyse the effect of this somewhat 
					# arbitrary choice of a parameter value for the model & analysis.
					# (Tau).

# Now set the model parameters that should be fitted (mutation.rate) and explored (virus.replication)

# Set the range of mutation rates, in units per bp per generation. (Mu)
mutation.rate.range = 10^(seq(-10, -5, by = 0.1))
num.mutation.rate = length(mutation.rate.range)

# Set the range of the virus replication parameter. A discreet parameter that specifies how descendents
# each genome has during the replication process. When = 1, you have a simple doubling model (i.e. 
# "bacterial replication"), when > 1 you go more and more towards a stamping machine model. (Rho)

# To speed things up, you can run different virus replication values in parallel. Note that you need to 
# then save the files with different names too (at the simulation loop).

virus.replication.range <- c(1, 3, 10)
num.virus.replication = length(virus.replication.range)	

#########################################################################################################

# Nested loops for generating model predictions over a range of model parameter values, to be used for 
# subsequent model fitting. Values to be varied include mutation rate and virus replication.

# Note: this part of the code can take very long to run. If you are running it with reasonable settings
# (E.g., max.stoch = 10^4, replicates = 1000, num.conditions [see below] > 50) it can easily take hours 
# when running for the bacmid, or several days for the whole genome.

# Generate a large array to store all data to be used for subsequent model fitting. The first two
# columns are the model parameter values, then a column for the frequency at which replicates
# contain mutated nucleotide positions (> detect.threshold). The highest value of mutations (i.e., 100)
# is arbitrary, so long as this value is not reached during the simulations. Additionally, arrays are made
# to store the results with alternative threshold values for detecting mutations. Finally there is counter
# to easily send the data to the correct row in the sim.data array.

setwd("C:/R_Data")
num.conditions = num.mutation.rate*num.virus.replication
sim.data <- array(NA, dim = c(num.conditions, (max.sim.data+3)))
colnames(sim.data) <- c("virus.replication", "mutation.rate", 0:max.sim.data)
sim.data.lo = sim.data
sim.data.hi = sim.data
counter.row = 1
print(num.conditions)

# Set random seed for reproducibility
set.seed(42)

for(f in 1:num.virus.replication) {
for(g in 1:num.mutation.rate) {

	# Set values of virus.replication and mutation.rate
	virus.replication = virus.replication.range[f]
	mutation.rate = mutation.rate.range[g]	
	
	# Empty vector to store the results for this set of conditions. Other arrays
	# for results with different mutation thresholds.
	mut.result <- rep(NA, replicates)
	mut.result.lo = mut.result
	mut.result.hi = mut.result
	
	for(h in 1:replicates) {
		  	
		# Generate the starting virus population, which is assumed to be clonal. 
		# Each position represents a locus (one bp), and its value represents the
		# frequency of a mutation at this locus.

		vir.pop = rep(0, genome.size)
		
		# Now run evolution on this genome.
			
		for(i in 1:number.passages) {
			
			# Evolve the virus population in 5 host insects.
			
			for(j in 1:5) {
				assign(paste0("vir.pop.", j), bottleneck.evolve(vir.pop, 
				bottleneck.size, final.size, virus.replication, mutation.rate))
			}
		
			# Determine the frequency of mutations in the pooled hosts, which serves 
			# as inoculum for the next round of infection.
			vir.pop = (vir.pop.1 + vir.pop.2 + vir.pop.3 + vir.pop.4 + vir.pop.5)/5
			
		} # end of i loop
  
		# Determine for which sites the mutation frequency is equal to or greater than the
		# threshold value, and send this result to array.
		mut.result[h] = length(which(vir.pop >= detect.thresholds[1]))
		mut.result.lo[h] = length(which(vir.pop >= detect.thresholds[2]))
		mut.result.hi[h] = length(which(vir.pop >= detect.thresholds[3]))
		
	} # end of h loop
 
	# Send the model parameter values to the results array
	sim.data[counter.row,c(1,2)] = c(virus.replication, mutation.rate) 
	sim.data.lo[counter.row,c(1,2)] = c(virus.replication, mutation.rate) 
	sim.data.hi[counter.row,c(1,2)] = c(virus.replication, mutation.rate) 
	
	# Determine the frequency of each possible outcome and send to table.
	for(k in 0:max.sim.data) {
		sim.data[counter.row, (k+3)] = length(which(mut.result == k))/replicates
		sim.data.lo[counter.row, (k+3)] = length(which(mut.result.lo == k))/replicates
		sim.data.hi[counter.row, (k+3)] = length(which(mut.result.hi == k))/replicates
	} # end of k loop
	
	# Save sim.data during each round, in case something goes wrong.			
	write.csv(sim.data,    file = "Baculo_mutation_simdata_BAC.csv")
	write.csv(sim.data.lo, file = "Baculo_mutation_simdata_BAC_lo.csv")
	write.csv(sim.data.hi, file = "Baculo_mutation_simdata_BAC_hi.csv")

	# Adjust the counter for rows in sim.data
	print(counter.row)
	counter.row = counter.row+1
	
} # end of g loop
} # end of f loop

#########################################################################################################

# Important: run the parameter settings for the simulations ("Basic model parameters that are not 
# explored.", block above between #####) prior to running the model fitting script below, 
# even if you have already completed the simulations.

# Next, based on the model predictions generated above we fit the model to the empirical data and
# hereby estimate the mutation rate. The model fitting only runs on the datasets for the 
# different thresholds at a time.

# Reintroduce some conditions set above in case the modelfitting is run in a new session.
num.conditions = num.mutation.rate*num.virus.replication

# Note: since this is approached is based on pseudolikelihoods, it can be beneficial for fitting to
# avoid frequency 0 predictions. These predictions can arise for model predictions because the 
# number of iterations cannot be very large. One quick solution would be to add a small value to
# each predicted frequency and normalize the data.

# First we have to exclude model predictions which exceed the treshold limit (i.e., those that reach
# a number of mutations higher than allowed by the results array). These conditions are removed 
# because whey will lead to problems with the likelihood estimation and are irrelevant (i.e., the
# value max.sim.data should be well above the observed number of mutations). Then determine how many
# conditions are left after pruning the results that exceed max.sim.data.
# Note, their are three different sets of simulation data (pru.sim.data) and experimental data
# (raw.data) with different values for the mutation detection threshold (0.5%, 1% and 2%). Make sure to
# always pair the correct ones when running the analysis!
setwd("C:/R_Data")
pru.sim.data <- read.csv(file = "Baculo_mutation_simdata_BAC.csv", header = TRUE) # 1% mutation frequency cutoff
# pru.sim.data <- read.csv(file = "Baculo_mutation_simdata_BAC_hi.csv", header = TRUE)  # 2% mutation cutoff
# pru.sim.data <- read.csv(file = "Baculo_mutation_simdata_BAC_lo.csv", header = TRUE)  # 0.5% mutation cutoff
pru.sim.data = data.matrix(frame = pru.sim.data[,-1]) 	# Delete extra column added during export
							# /import, and convert to array.
for(i in num.conditions:1) if(sum(pru.sim.data[i,3:max.sim.data]) != 1) pru.sim.data = pru.sim.data[-i,]
pru.conditions = nrow(pru.sim.data)

# Next we import the data. Here are some dummy data to work with for now. The format needed here is a
# vector with the number of mutations that were above detect.threshold. We then format the data to match
# the model results format (with counts instead of frequencies, however).
raw.data <- c(0, 0, 1, 1, 1) # 1% mutation frequency cutoff
# raw.data <- c(0, 0, 0, 0, 1) # 2% mutation cutoff
# raw.data <- c(2, 2, 1, 1, 2) # 0.5% mutation cutoff

emp.data <- rep(NA, (max.sim.data+1))
for(i in 1:(max.sim.data+1)) emp.data[i] = length(which(raw.data == (i-1)))

# Now we run a loop to determine the multinomial likelihood of each model prediction.
fit.results <- array(NA, dim = c(pru.conditions,3))
colnames(fit.results) <- c("virus.replication", "mutation.rate", "NLL")

for(i in 1:pru.conditions) {
	fit.results[i,1:2] = pru.sim.data[i,1:2]
	fit.results[i,3] = -dmultinom(x = emp.data, prob = as.numeric(pru.sim.data[i,3:(max.sim.data+3)]), log = TRUE)
}

# Write csv of complete output
write.csv(fit.results)

# Throw out cases where NLL = Inf
pru.fit.results = fit.results
for(i in pru.conditions:1) if(sum(pru.fit.results[i,3]) == Inf) pru.fit.results = pru.fit.results[-i,]

# Return best result
ord.nll = order(pru.fit.results[,3])
est.vir.rep = pru.fit.results[ord.nll[1],1]
est.mut.rate = pru.fit.results[ord.nll[1],2]

# Now make a plot of the results
dev.new(width= 5, height=3)
library(ggplot2)
gg.data <- as.data.frame(pru.fit.results)
gg.data$virus.replication <- as.factor(gg.data$virus.replication)
ggplot(data=gg.data, aes(x=log10(mutation.rate), y=NLL, linetype=factor(virus.replication), color=virus.replication)) + 
 geom_line(size = 1) + 
 xlab(" ") +  # Log10 [Mutation Rate] 
 ylab(" ") +  # NLL
 theme_classic() +
 scale_color_manual(values = c("violet", "blue", "green")) +
 scale_y_continuous(expand = c(0, 0), limits = c(0, NA)) +
 xlim(-10, -5)

###################

# Finally, we can bootstrap the data to determine a 95% confidence interval for the estimates.
num.bootstraps <- 1000
bootstrap.results <- array(NA, dim = c(num.bootstraps, 3))
colnames(bootstrap.results) <- c("virus.replication", "mutation.rate", "NLL")

# Set random seed for reproducibility
set.seed(42)

for(h in 1:num.bootstraps) {
	
	# Resample the data and reformat for multinomial function
	bs.data = sample(x = raw.data, size = length(raw.data), replace = TRUE)
	bs.emp.data <- rep(NA, (max.sim.data+1))
	for(i in 1:(max.sim.data+1)) bs.emp.data[i] = length(which(bs.data == (i-1)))
	
	# Loop to determine the multinomial likelihood of each model prediction for bs data.
	bs.fit.results <- array(NA, dim = c(pru.conditions,3))
		
	for(i in 1:pru.conditions) {
		bs.fit.results[i,1:2] = pru.sim.data[i,1:2]
		bs.fit.results[i,3] = -dmultinom(x = bs.emp.data, prob = pru.sim.data[i,3:(max.sim.data+3)], log = TRUE)
	}
	
	# Throw out cases where NLL = Inf, and then send the best result to bootstrap results array
	pru.bs.fit.results = bs.fit.results
	for(i in pru.conditions:1) if(sum(bs.fit.results[i,3]) == Inf) pru.bs.fit.results = pru.bs.fit.results[-i,]
	ord.nll = order(pru.bs.fit.results[,3])
	bootstrap.results[h,] = pru.bs.fit.results[ord.nll[1],]
	
	print(h)
	
} # end of h loop

# Now determine the CIs for both model parameters. Make an array to store the results and 
# add parameter estimates.
results.overview <- array(NA, dim = c(2,5))
colnames(results.overview) <- c("Lower_99_CI", "Lower_95_CI", "Parameter_Estimate", "Upper_95_CI", "Upper_99_CI")
rownames(results.overview) <- c("Virus_Replication", "Mutation_Rate")
results.overview[,3] = c(est.vir.rep, est.mut.rate)

# Determine which ordered results to take, order the data for each parameter and the populate 
# the results array.
ci.points = round(num.bootstraps*(c(0.005, 0.025, 0.975, 0.995)))
order.vr <- order(bootstrap.results[,1])
results.overview[1,c(1,2,4,5)] = bootstrap.results[order.vr[ci.points],1]
order.mr <- order(bootstrap.results[,2])
results.overview[2,c(1,2,4,5)] = bootstrap.results[order.mr[ci.points],2]

# Print the final result
write.csv(results.overview)
