## Supplementary material for "Empirical estimates of the mutation rate for an alphabaculovirus": Appendix 7 analysis script dN dS v1 AcMNPV.r.txt

##########################################################################################################
# Authors: Mark Zwart                                                                                    #
# Date: August 28, 2017                                                                                  #
# Modified: February 17, 2021                                                                            #
#                                                                                                        #
# This script is to determine dN/dS for the final clones of the AcMNPV mutation rate project, with 	 #
# experimental data generated by Ghulam Ali and subsequently analyzed by Dieke Boezen. First an		 #
# expectation is generated, and then compared to the data. This script is based on the script for the    #
# in vivo evolution project and its subsequent modification for the SpltNPV project.			 #
#                                                                                                        #
##########################################################################################################

setwd("C:/R_Data")
getwd()

# Load the AcMNPV bacmid with FB-dual and polyhedrin genome sequence. Here, I used a *.tsv of the coverage 
# for the bacmid, which contains the reference sequence but needs to be cleaned up (i.e., breaks due to 
# inserts from read mapping removed). Manual checks performed to make sure sequence is correct.

data.cov.bac  <- read.table("BAC_H752KDMXX_mapping_17022021.tsv", header = F)
ref.seq.a = as.vector(data.cov.bac[,4])
need.these = which(ref.seq.a == "A" | ref.seq.a == "C" | ref.seq.a == "T" | ref.seq.a == "G")
ref.seq = ref.seq.a[need.these]

# Load the legend for codons, the corresponding complementary codons should be included in columns 4 to 6.
# !!!NOTE!!! For simplicity the script evaluates the complementary sequence for 3' to 5' oriented genes,
# and does not use the reverse complementary sequence. Hereby you have been warned!

codon.legend <- read.csv("Codon_Legend.csv", header = F)
colnames(codon.legend) = c("1st_Base", "2nd_Base", "3rd_Base", "Comp_1st_Base", "Comp_2nd_Base", "Comp_3rd_Base") 


#########################################################################################################

# Read in the AcMNPV annotation (a modified CSV exported from CLC Genomics). This file contains
# only coordinates of the ORFs and no other genome features.

annotation.chrom <- read.csv("FastBacDual_polh_1805_annotation_for_dNdS.csv", head = T)

# You want to know three probabilities: that a mutation is intergenic, synonymous or non-synonymous for 
# the whole genome. The simplist is the probability of an intergenic mutation, since every
# mutation outside an ORF meets this criterion. However, due to mutational biases and the need to compare
# to the occurence of S mutations, you want to know how many of each base there are in the intergenic 
# spaces. Let's start with this then. Here we perform this analysis only for the native AcMNPV intergenic
# regions, ignoring the bacmid insert.

# Make a huge vector which contains the position of all nucleotides within ORFs

coding.positions = vector('numeric')
dim.chrom <- dim(annotation.chrom)
genes.chrom <- dim.chrom[1]

for(i in 1:genes.chrom) coding.positions = c(coding.positions, (annotation.chrom[i,5]:annotation.chrom[i,6]))

# Here we tack the bacmid regions onto the coding regions, so they are not considered an intergenic
# region here.
non.coding.bacmid <- c(4411:6621, 7460:16920) # 6622:7459 is the inserted polh copy don't duplicate it here.
coding.positions.plusbac = c(coding.positions, non.coding.bacmid) 

# Next, parse the nucleotide data for all other positions from the genome data.
intergenic.seq = ref.seq[-coding.positions.plusbac]
intergenic.length = length(intergenic.seq)
intergenic.length

# Now, tally the bases for the intergenic sequences. Generate an array to store the results, then determine
# the absolute and relative frequencies.

intergenic.results = array(NA, dim = c(4,2))
colnames(intergenic.results) <- c("Count", "Frequency")
rownames(intergenic.results) <- c("A", "C", "G", "T")

intergenic.results[1,1] = length(which(intergenic.seq == "A" ))
intergenic.results[2,1] = length(which(intergenic.seq == "C" ))
intergenic.results[3,1] = length(which(intergenic.seq == "G" ))
intergenic.results[4,1] = length(which(intergenic.seq == "T" ))
intergenic.results[,2] = intergenic.results[,1]/intergenic.length

write.csv(intergenic.results, file = "Intergenic_Results_AcMNPV.csv")

# We can do the same for the bacmid region, just as a control:

nc.bacmid.seq = ref.seq[non.coding.bacmid]
nc.bacmid.length = length(nc.bacmid.seq)
nc.bacmid.length

nc.bacmid.results = array(NA, dim = c(4,2))
colnames(nc.bacmid.results) <- c("Count", "Frequency")
rownames(nc.bacmid.results) <- c("A", "C", "G", "T")

nc.bacmid.results[1,1] = length(which(nc.bacmid.seq == "A" ))
nc.bacmid.results[2,1] = length(which(nc.bacmid.seq == "C" ))
nc.bacmid.results[3,1] = length(which(nc.bacmid.seq == "G" ))
nc.bacmid.results[4,1] = length(which(nc.bacmid.seq == "T" ))
nc.bacmid.results[,2] = nc.bacmid.results[,1]/nc.bacmid.length

write.csv(nc.bacmid.results, file = "Noncoding_Bacmid_Results_AcMNPV.csv")

# For other analyses performed later (mutation spectrum), we also want to know the distribution 
# of nucleotides over the whole genome.

ref.seq.length = length(ref.seq)
ref.seq.length

ref.seq.results = array(NA, dim = c(4,2))
colnames(ref.seq.results) <- c("Count", "Frequency")
rownames(ref.seq.results) <- c("A", "C", "G", "T")

ref.seq.results[1,1] = length(which(ref.seq == "A" ))
ref.seq.results[2,1] = length(which(ref.seq == "C" ))
ref.seq.results[3,1] = length(which(ref.seq == "G" ))
ref.seq.results[4,1] = length(which(ref.seq == "T" ))
ref.seq.results[,2] = ref.seq.results[,1]/ref.seq.length

write.csv(ref.seq.results)

#########################################################################################################

# Now comes the more complicated part: you want to know how often every codon is present in the complete 
# coding part of the genome. I will do this gene by gene, and then codon for codon with genes. Given that
# this script is computationally very ineffecient, and the size of the coding genome, this part of the 
# script probably runs very slowly. The codon.legend is uploaded as a csv, and also used to make a results 
# array. For the AcMNPV this takes about 10 minutes on my PC, so not too bad.

codon.count = rep(0, 64)
codon.freq  = rep(NA,64)
coding.results = cbind(codon.legend, codon.count, codon.freq)

start.codon = c("A", "T", "G")
anti.start.codon = c("T", "A", "C") 

for(i in 1:genes.chrom) {
	
	# Determine the characteristics of the gene of interest, absolute value just in case
	
	five.prime.end = annotation.chrom[i,5]
	three.prime.end = annotation.chrom[i,6]
	gene.length = abs(five.prime.end - three.prime.end)
	
	# For genes with a 5'to 3'sense orientation, analyze here
	
	if(sum(start.codon == c(ref.seq[five.prime.end], ref.seq[five.prime.end+1], ref.seq[five.prime.end+2])) == 3 ) {
		
		triplet.now = five.prime.end
				
		while(triplet.now < three.prime.end) {
			
			# Here the script identifies which codon is present at a position, and 
			# tallies the counts.
			
			codon.now = c(ref.seq[triplet.now], ref.seq[triplet.now+1], ref.seq[triplet.now+2])
			
			for(j in 1:64) {
				ref.codon = codon.legend[j,(1:3)]
				if(sum(codon.now == ref.codon) == 3) {
					coding.results[j,7] = coding.results[j,7] + 1
					} # end of if statment
			} # end of j loop
			
			triplet.now = triplet.now + 3
			
		} # end of while loop
	} # end of first if loop
		
	# Second if for 3' to 5' oriented genes here.

	if(sum(anti.start.codon == c(ref.seq[three.prime.end], ref.seq[three.prime.end-1], ref.seq[three.prime.end-2])) == 3 ) {

		triplet.now = three.prime.end
		
		while(triplet.now > five.prime.end) {
			
			# Here the script identifies which codon is present at a position, and 
			# tallies the counts.
			
			codon.now = c(ref.seq[triplet.now], ref.seq[triplet.now-1], ref.seq[triplet.now-2])
			
			for(j in 1:64) {
				ref.codon = codon.legend[j,(4:6)]
				if(sum(codon.now == ref.codon) == 3) {
					coding.results[j,7] = coding.results[j,7] + 1
					} # end of if statment
			} # end of j loop
			
			triplet.now = triplet.now - 3
			
		} # end of while loop
	} # end of first if loop
	
	print(i)
	
} # end of i loop

coding.results[,8] = coding.results[,7]/sum(coding.results[,7])
write.csv(coding.results, file = "Coding_Results_AcMNPV.csv")


#########################################################################################################
#                                                                                                       #
# Here the acutal dN/dS calculations are made, after generating predictions for chrom and plasmid.      #
#                                                                                                       #
# Update: in the modified script, only the expectations for N and S are made here, final calculations   #
# need to be done post hoc in an Excel sheet.                                                           #
#                                                                                                       #
# Update 2:  in the absence of relevant data on the mutational spectrum or biases in DNA viruses, I     #
# have kept the analysis and assumed no mutational biases occur.                                        #
#													#
# Update AcMNPV: it feels awkward to estimate a mutation bias and dN/dS from the same data, so I simply #
# assume there is no mutation bias. 									#
#                                                                                                       #
#########################################################################################################

# Next, we want to use the information in the two previously generated arrays to get an expectation for 
# the occurence of intergenic, synonymous and non-synonymous mutations.
# Load the codon legend and the mutation spectrum information. 
# Also generate an array of intergenic and coding results combining chromosome and plasmid data. (*.whole)

# The script can be run from here, since all previous information generated has been saved to *.csv.
# Note: code related to plasmids again removed, since it is irrelevant here.

intergenic.results.whole <- read.csv("Intergenic_Results_AcMNPV.csv", header = TRUE)
coding.results.whole <- read.csv("Coding_Results_AcMNPV.csv", header = TRUE)

# For each codon, you want to know what the frequency of synonymous and nonsynonymous subsitutions is,
# corrected for the mutational biases as given in the mutational spectrum data. Generate an array to
# store the final data, and then run a loop where you calculate the frequency that a mutation will be 
# nonsynonymous or synonymous. Load the necessary data to do so, including codon legend, the
# translation index (codon legend + AA coded for) and emperical mutation spectrum data. Since we do 
# not have such data for DNA viruses, we assume no mutational biases.

codon.legend <- read.csv("Codon_Legend.csv", header = FALSE)
translation.index <- read.csv("Codon_Legend_Expanded.csv", header = TRUE)
mutation.spectrum <- read.csv("Mutational_spectrum_data_NULL.csv", header = TRUE)

codon.s.ns = cbind(codon.legend[,1:3], Nonsyn = rep(NA, 64), Syn = rep(NA, 64))

for(i in 1:64) {
	
	# Set reference codon (i.e., ancestral sequence)
	# Set reference aminoacid. This can be done straight from i because the translation index
	# organisation is identical.
	
	ref.codon = codon.s.ns[i, c(1:3)]
	ref.aa = translation.index[i,5]
		
	# Make an array to store the results for this codon, and then test the effects of all possible 
	# mutations in this codon.
	
	results.this.codon = array(NA, dim = c(9, 3))
	colnames(results.this.codon) <- c("Mod_Base", "Prob_Mut", "NS_or_S")
		
	for(j in 1:3) {
		
		# Determine what the base now is, and which mutations should be considered. Base.now
		# is assign separately because using "base.now = ref.codon[j]" leads to problems with
		# logical statements. 
		
		first.row = (3*(j-1))+1
		last.row  = (3*(j-1))+3
		if(ref.codon[j] == "A") {
			results.this.codon[c(first.row:last.row), 1] = c("C", "G", "T")
			base.now = "A"
		}
		if(ref.codon[j] == "C") {
			results.this.codon[c(first.row:last.row), 1] = c("A", "G", "T")
			base.now = "C"
		}	
		if(ref.codon[j] == "G") {
			results.this.codon[c(first.row:last.row), 1] = c("A", "C", "T")
			base.now = "G"
		}
		if(ref.codon[j] == "T") {
			results.this.codon[c(first.row:last.row), 1] = c("A", "C", "G")
			base.now = "T"
		}
		
		# Now evaluate the frequency of each possible change at the jth position of the triplet,
		# and whether each mutation is NS or S.
		
		for(k in 1:3) {
			
			# Make the codon to be compared to the reference (test.codon)
			
			test.codon = ref.codon
			row.now = (3*(j-1))+k
			test.codon[j] = results.this.codon[row.now,1]
								
			# Record the probability of a particular mutation
			
			base.change = which(mutation.spectrum[,1] == results.this.codon[row.now,1] & mutation.spectrum[,2] == base.now)
			results.this.codon[row.now,2] = mutation.spectrum[base.change,4]
			
			# Determine which AA corresponds with the test codon
			
			for(l in 1:64) {
				is.codon = codon.legend[l,(1:3)]
				if(sum(test.codon == is.codon) == 3) test.aa = translation.index[l,5]
			} # end of j loop
			
			# Determine whether nonsynonymous and store result in array
			
			results.this.codon[row.now,3] = (ref.aa != test.aa)
						
		} # end of k loop
				
	} # end of j loop
	
	ns.changes.now = which(results.this.codon[,3] == TRUE)
	codon.s.ns[i,4] = sum(as.numeric(results.this.codon[ns.changes.now,2]))
	codon.s.ns[i,5] = sum(as.numeric(results.this.codon[-ns.changes.now,2]))
	
	print(i)
	
} # end of i loop

codon.s.ns <- cbind(codon.s.ns, Freq_NS_per_AA = (codon.s.ns[,4]/(codon.s.ns[,4]+codon.s.ns[,5])))
hist(codon.s.ns[,6], n = 20, xlim = c(0,1), col = "blue", main = "Histogram of NS mutation probability per codon", xlab = "Frequency of NS mutations")


#########################################################################################################

# Now generate predictions for the number of synonymous, nonsynonymous, and intergenic mutants expected
# from the whole AcMNPV reference sequence.

expected.ns = sum(coding.results.whole[,8]*codon.s.ns[,4])
expected.ns

expected.s  = sum(coding.results.whole[,8]*codon.s.ns[,5])
expected.s

# For expected intergenics, consider the frequency of all mutations which can occur for a given base, 
# taking into account mutational bias.

correction.intergenic = numeric(4)
correction.intergenic[1] = sum(mutation.spectrum[which(mutation.spectrum[,1] == "A"),4])
correction.intergenic[2] = sum(mutation.spectrum[which(mutation.spectrum[,1] == "C"),4])
correction.intergenic[3] = sum(mutation.spectrum[which(mutation.spectrum[,1] == "G"),4])
correction.intergenic[4] = sum(mutation.spectrum[which(mutation.spectrum[,1] == "T"),4])
expected.i  = sum(intergenic.results.whole[,2]*correction.intergenic)

expected.i

# Subsequent analysis is all done in an Excel sheet for simplicity.
