## Supplementary material for "Empirical estimates of the mutation rate for an alphabaculovirus": Appendix 8 Confidence intervals dNdS by bootstrapping.txt

##########################################################################################################
# Authors: Mark Zwart                                                                                    #
# Date: June 2, 2021											 #
#                                                                                                        #
# Using bootstrapping to determine 95% fiducial interval for dN/dS					 #
#                                                                                                        #
##########################################################################################################

# Data: from the baculovirus mutation rate project from Dieke and Ghulam, only the data with the lowest
# threshold for mutation detection (0.5%) appear to be useful, as in all other cases the number of 
# mutations is too low.
reps  <- c(  1,  2,  3,  4,  5)  # the replicate lineages A to E
n.mut <- c(  3,  3,  1,  3,  3)  # Non-synonymous mutations per lineage
s.mut <- c(  3,  2,  2,  2,  1)  # Synonymous mutations per linage
b.mut <- c(  2,  2,  1,  1,  2)  # Intergenic mutations in the bacmid region only
num.obs <- length(reps)

# Nul expectations as calculated from genome: the number of possible mutations in each category. See other
# R script to generate these numbers.
n.nul <- 297236
s.nul <-  76381 
b.nul <-  35016

# Just as a check: dN/dS calculation
p.n = sum(n.mut)/n.nul
p.s = sum(s.mut)/s.nul
p.b = sum(b.mut)/b.nul
d.n = (3/4)*log(1 - (4*p.n/3))
d.s = (3/4)*log(1 - (4*p.s/3))
d.b = (3/4)*log(1 - (4*p.b/3))
d.n/d.s
d.b/d.s


# Make an array to store the bootstrap results
bs.num <- 1000 # number of bootstraps
bs.data <- array(NA, dim = c(bs.num, 2))
colnames(bs.data) <- c("dN/dS", "dI_bac/dS")

# Loop for bootstrapping
for(i in 1:bs.num) {
	
	hit.me = sample(x = (1:num.obs), size = num.obs, replace = TRUE)
	bs.n.mut = n.mut[hit.me]
	bs.s.mut = s.mut[hit.me]
	bs.b.mut = b.mut[hit.me]
	
	# dN/dS calculation & send result to array
	p.n = sum(bs.n.mut)/n.nul
	p.s = sum(bs.s.mut)/s.nul
	p.b = sum(bs.b.mut)/b.nul
	d.n = (3/4)*log(1 - (4*p.n/3))
	d.s = (3/4)*log(1 - (4*p.s/3))
	d.b = (3/4)*log(1 - (4*p.b/3))
	bs.data[i,] = c(d.n/d.s, d.b/d.s)	
	
} # end of i loop

# Determine 95% CIs
limits = c(25, 975)
ord.dn.ds = order(bs.data[,1])
for(i in 1:2) print(bs.data[ord.dn.ds[limits[i]],1])
ord.di.ds = order(bs.data[,2])
for(i in 1:2) print(bs.data[ord.di.ds[limits[i]],2])


#############################

# Finally, run a one-sample t test to look for significance against a test value of 1
p.n = n.mut/n.nul
p.s = s.mut/s.nul
p.b = b.mut/b.nul
d.n = (3/4)*log(1 - (4*p.n/3))
d.s = (3/4)*log(1 - (4*p.s/3))
d.b = (3/4)*log(1 - (4*p.b/3))
dn.ds = d.n/d.s 
db.ds = d.b/d.s 
t.test(x = dn.ds, mu = 1)
t.test(x = db.ds, mu = 1)
